# Functional Mapping of the *Trypanosoma cruzi* Serinome by Fluorophosphonate Activity-Based Protein Profiling

**DOI:** 10.64898/2026.07.18.739127

**Authors:** Jaime A. Isern, María Gabriela Mediavilla, Exequiel O.J. Porta, Marcelo L. Merli, María Sol Ballari, Julia A. Cricco, Guillermo R. Labadie, Patrick G. Steel

## Abstract

Serine hydrolases (SHs) constitute one of the largest enzyme superfamilies in eukaryotes, yet their roles in *Trypanosoma cruzi,* the causative agent of Chagas disease, remain largely uncharacterized. Here, we report an activity-based chemoproteomic map of the *T. cruzi* epimastigote serinome by combining genome-informed *in silico* curation with whole-cell activity-based protein profiling (ABPP) using a panel of cell-permeable fluorophosphonate (FP)-alkyne probes. Whole-cell labelling followed by label-free quantitative proteomics (LFQ-MS), identified 37 enriched SH-like proteins, including 35 with conserved or partially conserved catalytic triad/dyad features, spanning lipases, peptidases, esterases, and previously uncharacterized hydrolases. The 35 SHs represent approximately 63 % of the 56 predicted SHs retained after catalytic-site curation. Domain architecture analysis revealed broad structural diversity, while orthologue-based localization data suggested association with multiple subcellular compartments, including glycosomal, mitochondrial, and endosomal localizations. Gene Ontology enrichment highlighted lipid metabolic and catabolic processes as dominant functional themes, and protein-protein interaction network analysis supported functional connectivity among the captured enzymes. Several identified SHs, including oligopeptidase B, prolyl oligopeptidase Tc80, serine carboxypeptidase CPB1, and phospholipase A1 (PLA1) have previously been characterized in trypanosomatids as virulence factors and as mediator of host-pathogen interactions. Together, these findings establish a fluorophosphonate-based chemoproteomic resource for the kinetoplastid community and prioritize probe-accessible active *T. cruzi* SHs for future functional validation and antiparasitic inhibitor discovery.

## Introduction

Chagas disease, caused by the protozoan parasite *T. cruzi*, is a life-threatening neglected tropical disease (NTD) endemic to Latin America and increasingly recognized as a global health concern due to migration and climate change. It is estimated that between 6 and 8 million people are currently infected, with more than 75 million people considered at risk of infection across 21 endemic countries.^1,2^ Transmission occurs primarily through contact with the faeces of triatomine insects, but can also occur congenitally, via blood transfusion, organ transplantation, or orally through contaminated food or drink.^2,3^ The disease progresses from an often-asymptomatic acute phase to a chronic phase, which may lead to irreversible cardiac, gastrointestinal, and neurological complications. If left untreated, chronic Chagas disease can result in severe disability and death, as evidenced by approximately 10,000 deaths attributed to the disease each year.^1,4^

Despite its significant disease burden, Chagas disease remains severely underdiagnosed and undertreated. Current therapeutic options are limited to two nitro-heterocyclic drugs, benznidazole and nifurtimox, both developed over 50 years ago. These drugs are primarily effective during the acute phase, show limited efficacy in chronic infections, and are associated with substantial side effects and treatment discontinuation rates.^5,6^ Moreover, no vaccine is currently available. These limitations underscore the urgent need to identify new drug targets and therapeutic strategies, in line with the World Health Organization’s 2021–2030 roadmap for NTDs.^7^

The serine hydrolase (SH) superfamily constitutes one of the largest and most functionally diverse classes of enzymes, encompassing proteases, esterases, and lipases. In *T. cruzi*, proteases represent key virulence factors involved in immune evasion, nutrient acquisition, and host tissue invasion.^8,9^ The cysteine protease family has been extensively studied in *T. cruzi*, identifying vital proteases such as cruzipain, involved in cell differentiation, metabolism, invasion of the host cell and evasion of the immune response.^10–14^ In contrast, comparatively little is known about the role of SHs in *T. cruzi* biology. In other protozoan parasites such as *Leishmania* spp., SHs have been shown to play pivotal roles in virulence, stage differentiation and host-pathogen interactions.^15–20^ These findings suggest that SHs may also be critical to the *T. cruzi* life cycle and pathogenicity and could offer underexplored opportunities for therapeutic intervention.

Although omics approaches provide valuable information on transcript or protein abundance, abundance does not necessarily correlate with functional activity. Functional annotation of enzyme families such as SHs therefore requires approaches that give insights into enzyme activity. In this context, activity-based protein profiling (ABPP) has emerged as a powerful chemoproteomics technique for interrogating enzyme function in complex biological systems. ABPP utilizes mechanism-based probes that covalently bind to active-site residues, enabling the selective detection of active enzymes within native proteomes against their non-active zymogens.^21–23^ This technology has been particularly transformative for the functional annotation of serine hydrolases, with fluorophosphonate (FP) probes providing a robust platform to interrogate catalytically active members of this enzyme family in complex proteomes.^17,24,25^ Structural modifications to the FP scaffold can further enhance probe selectivity and cell permeability, providing valuable tools for enzyme profiling in both host and pathogen contexts.

We have recently described the development and application of a series of cell-permeable fluorophosphonate (FP)-based probes to characterize the active serinome of *Leishmania* spp. These proved effective in comparing serinomes across different *Leishmania* species and life-cycle stages, including the disease-critical intramacrophage amastigote, providing insights into the host–pathogen interactome. Here, we further demonstrate their effectiveness for functional profiling kinetoplastid parasites,^17^ applying the probes to profile the active serinome of *T. cruzi*. To our knowledge, the active serine hydrolases of *T. cruzi* have not previously been profiled directly. Here we provide a functional analysis of SHs in *T. cruzi*, combining ABPP with genome wide *in silico* curation to establish a valuable activity-based chemoproteomic resource for the *T. cruzi* serinome that identifies a prioritized set of catalytically active SHs for future functional validation and inhibitor discovery.

## Results and Discussion

### *In silico* prediction of the *T. cruzi* serinome

To establish a comprehensive reference framework for the *T. cruzi* serinome, we first performed a genome-wide *in silico* survey of predicted SHs using the Dm28c 2018 genome reported by Berná *et al.*^26^ (**Fig. 1**). A total of 135 proteins were identified as putative SHs, characterized by the presence of serine hydrolase domains annotated in Pfam, InterPro, CDD, or Panther, or by alignment to the peptidase database MEROPS. Following clustering to remove redundancy and discarding protein fragments, 73 unique proteins remained as putative SHs (**Table 1** and **Supplementary Data 1**). These were classified as esterases, serine peptidases, amidases, or unclassified SHs according to the identified domains or additional information shown in Supplementary Data 1.

To refine this candidate set, we assessed whether each protein possessed catalytic features consistent with serine hydrolase activity. Sequence alignments against reference proteins with known catalytic residues were combined with inspection of AlphaFold-predicted structures to determine whether catalytic residues were positioned appropriately in three-dimensional space (**Supplementary Fig. S1**). For canonical SHs, we assessed the presence of an S-H-D/E catalytic triad, with the nucleophilic serine typically located within a conserved GxSxG motif. Catalytic geometry was scored as conserved when the Ser O_γ_–His N_ε_2 and His–Asp/Glu interatomic distances were within 5 Å in the AlphaFold model and the participating residues had an average pLDDT ≥ 80, with selected borderline cases accepted when pLDDT values were between 60 and 79. For amidase-signature enzymes a Ser–Ser–Lys constellation was required, and for rhomboids a Ser–His dyad. Borderline cases were classified as partially conserved, with per-protein metrics reported in **Supplementary Data 1**. Representative active-site structures are shown in **Fig. 1b**. Ultimately, of the 73 putative SHs, 56 gene products were retained as SH candidates based on conserved or partially conserved catalytic triad/dyad features in *T. cruzi* strain Dm28c (**Supplementary Fig. S1**).

Of the 56 total genes identified, the majority encoded esterases (29 genes), followed by serine peptidases (19 genes), unclassified SHs (7 genes), and one amidase. Most serine peptidase families (S1, S8, S9, S26, S28, S33, S51, and S54) contained one to four candidate genes, with each gene generally present as a single copy in the genome. Notably, peptidase S15 and S10 showed copy-number expansion, with three and twelve predicted copies respectively. Among the esterases, most classes were represented by only 1–2 genes or copies. In contrast, GDSL lipases and lipase class 3 showed notably higher representation. For example, GDSL lipase C4B63_184g21 had 3 copies, while lipase class 3 comprised 10 unique genes, three of which had 2 copies each, and C4B63_104g97 alone accounted for 17 copies (**Table 1** and **Supplementary Data 1).**

**Figure 1.**
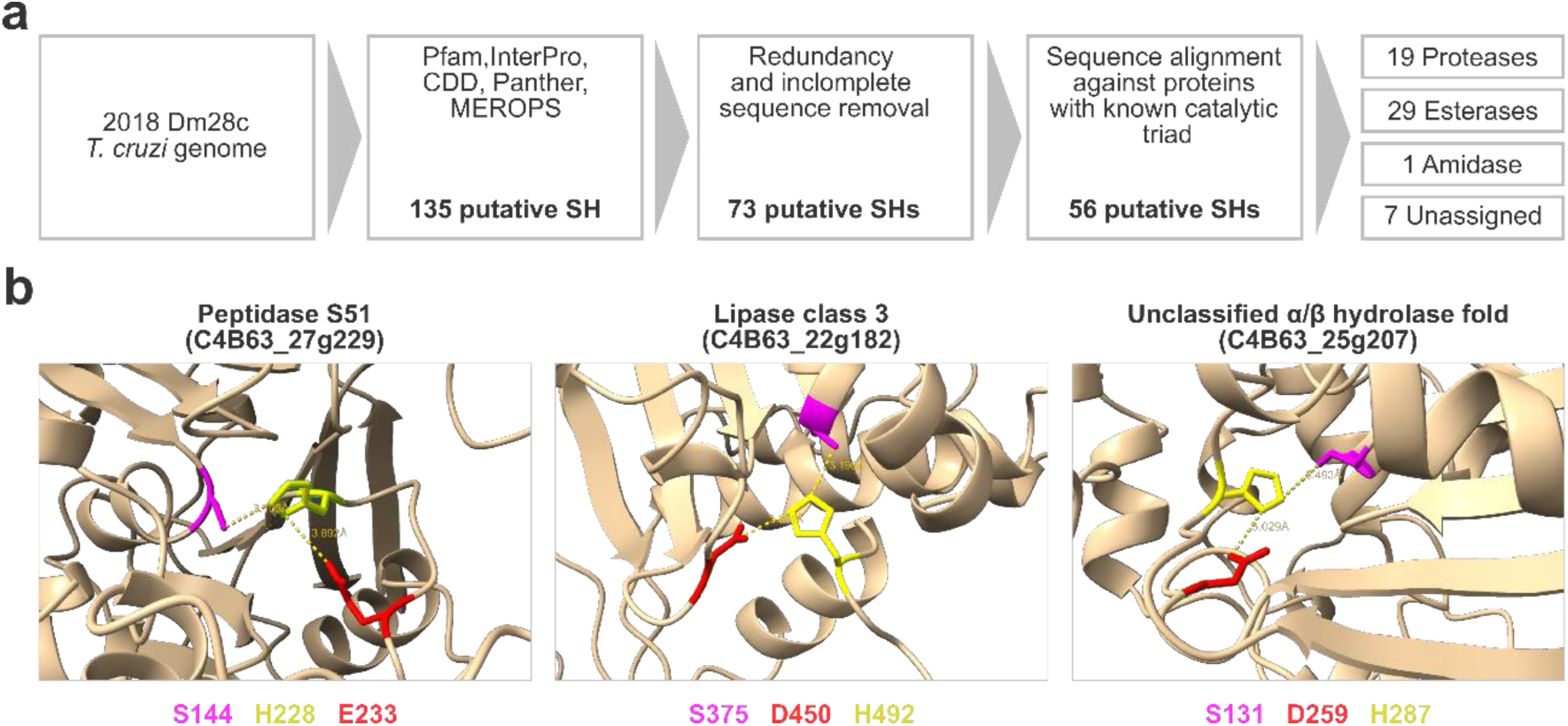
**a.** *In silico* serinome workflow. **b**. Close-up view of the active site showing the spatial arrangement of the catalytic S–H–D/E triad in representative proteins from the full dataset (Peptidase S51, Lipase class 3, and an unclassified hydrolase). The pLDDT values are 98.25 (S144), 97.88 (H228), and 89.19 (E233) for Peptidase S51; 86.44 (S375), 63.22 (H492), and 69.44 (D450) for Lipase class 3; and 91.12 (S131), 89.88 (H287), and 89.88 (D259) for the unclassified hydrolase. Residue numbers are indicated in the color-coded legend, and dashed lines denote interatomic distances (Å). AlphaFold-predicted structures (AF-A0A2V2VFP5-F1-model_v6, AF-A0A2V2UKM7-F1-model_v6, and AF-A0A2V2VHL2-F1-model_v6) were obtained from the AlphaFold Protein Structure Database.

**Table 1.** Serine Hydrolases identified *in silico* in the *T. cruzi* genome. We identified 73 accession numbers that correspond to putative SHs and verified that each sequence has the catalytic triad or dyad with the nucleophilic serine. The table shows the list of accession numbers classified as esterase, serine peptidase, or amidases according to Pfam, product description, MEROPS classification, other identified domains, or catalytic triad (**Supplementary Data 1**). The number of copies is shown in parentheses after the accession number for any value greater than one. A single representative accession number is reported for multi-copy genes. Fifty six out of the 73 genes were selected based on the presence of conserved or partially conserved catalytic triad. *The catalytic triad is partially conserved.

| Activity Classification |  | Catalytic Triad/Dyad |  |
| --- | --- | --- | --- |
| fold | Family or Classification | Conserved | Not conserved |
| Serine Peptidases |  |  |  |
| β-barrel core fold | S1 | C4B63_23g231 C4B63_9g401 | C4B63_17g44<br>C4B63_75g55<br>C4B63_48g166 |
|  | S26 | C4B63_31g105<br>C4B63_13g150 |  |
| subtilisin-type 3-layer $\alpha/\beta$ sandwich fold | S8 | C4B63_10g296 C4B63_95g13 | |
| *β-propeller fold and α/β hydrolase fold* | S9 | C4B63_101g15 C4B63_63g37<br>C4B63_6g564 C4B63_122g16 |  |
| *α/β hydrolase fold* | S10 | C4B63_168g38 (12) |  |
|  | S15 | C4B63_6g235 (3) |  |
|  | S28 | C4B63_57g112<br>C4B63_17g176* |  |
|  | S33 | C4B63_2g814 C4B63_84g83<br>C4B63_2g162 |  |
| *trpG-type fold* | S51 | C4B63_27g229 |  |
| *rhomboid fold* | S54 | C4B63_34g300 | C4B63_38g78 |
| *GAIN fold* | S59 |  | C4B63_19g195 |
| **Amidases** |  |  |  |
| *amidase signature fold* | Fatty-acid amide hydrolase (FAAH) | C4B63_55g66 |  |
| **Esterases** |  |  |  |
| *patatin fold* | Patatin-like Lipase | C4B63_18g341 C4B63_7g246 | C4B63_95g29<br>C4B63_7g337 |
| *SGNH hydrolase fold* | GDSL Lipase | C4B63_184g21 (3)<br>C4B63_13g128 |  |
| *α/β hydrolase fold* | Lipase class 3 | C4B63_25g256<br>C4B63_22g182 (2)<br>C4B63_104g97 (17)<br>C4B63_10g143 C4B63_171g8<br>(2) C4B63_10g145<br>C4B63_20g182 C4B63_4g367<br>C4B63_20g60 C4B63_10g276<br>(2)* | C4B63_2g44 |
|  | Phospholipid: diacylglycerol acyltransferase (PDAT) | C4B63_5g420 (2) |  |
|  | Phospholipase A | C4B63_2g791 | C4B63_2g790 |
|  | GPI inositol-deacylase PGAP1 | C4B63_23g274 C4B63_2g218* |  |
|  | LID Hydrolase | C4B63_8g548 |  |
|  | Phospholipase/ Carboxylesterase | C4B63_32g268<br>C4B63_14g203 |  |
|  | Bifunctional carbohydrate esterase (CE) | C4B63_83g76 |  |
|  | Unclassified | C4B63_44g168 (3)<br>C4B63_232g4 (2)<br>C4B63_46g85 C4B63_17g73<br>C4B63_6g260 C4B63_35g372*<br>TCDM_11513(5) |  |
| Unclassified |  |  |  |
| $\alpha/\beta$-Hydrolase fold | | C4B63_20g152<br>C4B63_42g215<br>C4B63_60g130<br>C4B63_12g197<br>C4B63_25g206<br>C4B63_25g207 C4B63_48g32<br>(2) | C4B63_25g205<br>C4B63_14g216<br>C4B63_7g399<br>C4B63_14g209<br>C4B63_44g234<br>C4B63_20g186<br>C4B63_24g264 (2)<br>C4B63_25g137 (2) |

### Chemoproteomic Profiling of Serine Hydrolases

Bioinformatics-driven approaches have become indispensable for predicting protein function, pathway integration, and druggability potential. Comparative genomics, sequence-based classification, structural modelling, and machine learning algorithms enable the identification of conserved domains, essentiality, and active site features that help prioritize enzymes for further biological characterization and therapeutic exploration. Databases such as TDR Targets and TriTrypDB further integrate functional genomics with chemical biology information, facilitating target prioritization based on genetic essentiality, lack of human homologues, and druggability indices.^27,28^ For enzyme families such as the SHs, this computational framework is invaluable for distinguishing catalytically active members, identifying non-canonical variants, and informing probe or inhibitor design. However, these methods do not report on activity, a particular limitation for SHs, as many proteases are initially produced as zymogens and only activated in response to specific signals. This provides a strong rationale for ABPP, which can identify catalytically active enzymes directly in a biological context.

We therefore used a panel of cell-permeable FP probes to experimentally profile the active SH landscape of *T. cruzi*. In previous work we had reported the synthesis of a set of alkyl-, benzyl- and aryl fluorophosphonate activity-based probes (ABPs) (**Fig. 2**) and their application to examine SHs in *Leishmania*.^17,29^ Initial studies explored the use of these probes with *T. cruzi* cell lysate. These extracts were prone to agglutination, likely associated with the complex *T. cruzi* surface glycocalyx,^30–34^ which manifested as poorly resolved, aggregated material on SDS-PAGE. Although this could be partially mitigated, no labelling was observed, suggesting that agglutination was not the sole barrier and that access of the ABPs to their targets in the lysate may have been also limited. Consequently, exploiting the cell-permeable nature of the probes, a whole-cell labelling approach was adopted. Importantly, as demonstrated in our previous studies,^17^ this minimizes target loss associated with lysis.

**Figure 2.**
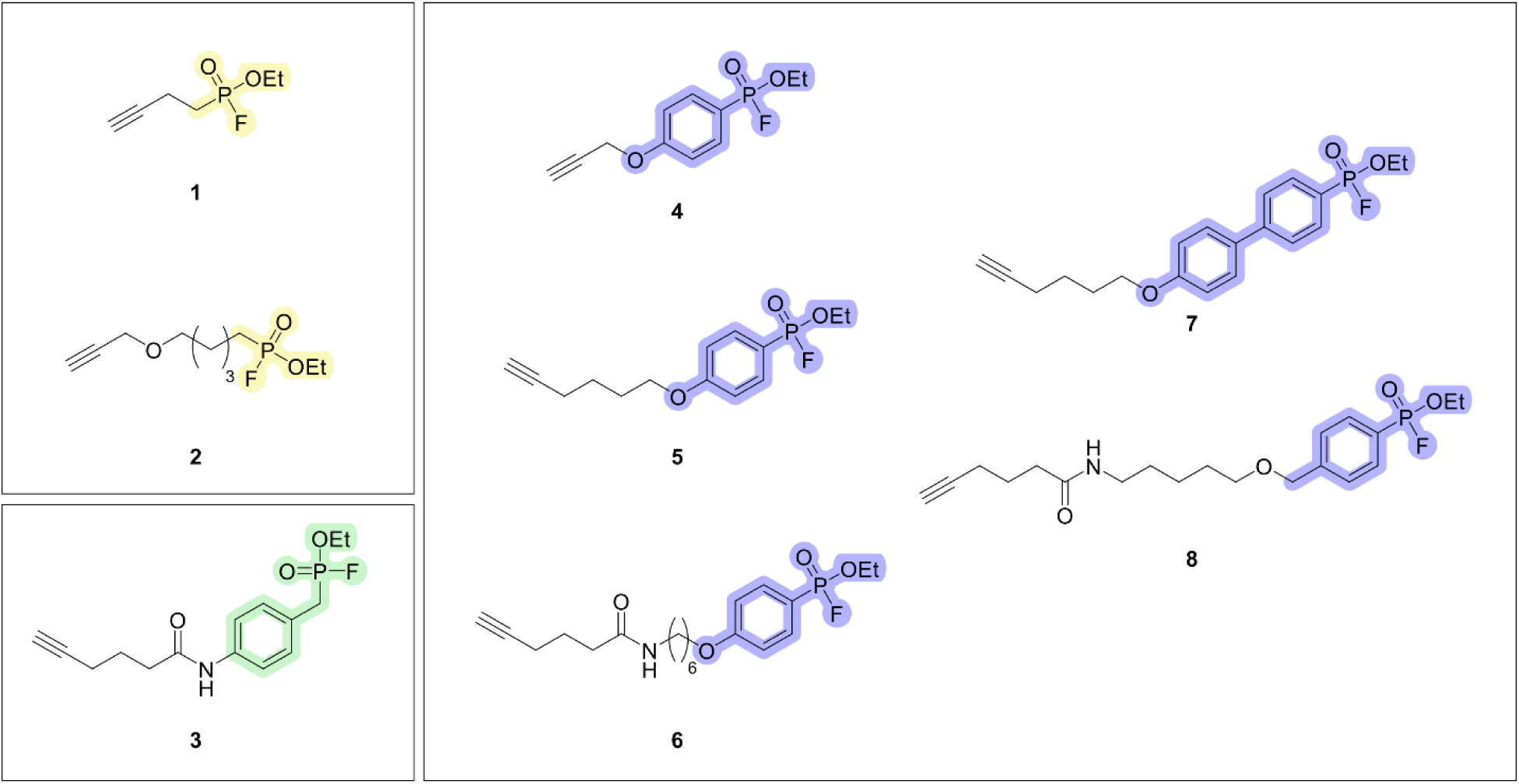
Structures of the cell-permeable activity-based fluorophosphonate (FP-alkyne) probes used in this study: alkyl-FPs (yellow), benzyl-FP (green), and aryl-FPs (purple).

To identify probe-labelled SHs in *T. cruzi*, we employed a label-free quantitative proteomics workflow. After optimisation of the lysis conditions, live parasites were incubated with FP-alkyne probes, followed by cell lysis, click chemistry with biotin-N_3_, streptavidin enrichment, reduction, alkylation, digestion, and LC-MS/MS analysis. Three biological replicates were analysed per condition. Differential protein abundance between probe-treated and control samples was assessed using empirical Bayes moderated t-tests implemented in the limma package.^35^ Proteins with a log_2_ fold-change > 1 and a p-value < 0.05 were considered significantly enriched. Benjamini–Hochberg-adjusted values were computed but not used for thresholding, to avoid discarding genuine low-abundance hydrolases in this discovery-oriented workflow, and are reported per protein in **Supplementary Data 2-3**.

### Genome assembly influences serine hydrolase identification

Accurate identification of the complete repertoire of enriched serine hydrolases in *T. cruzi* depends on the quality and completeness of the reference genome used for protein annotation. However, the availability of high-quality whole-genome assemblies for *T. cruzi* remains limited due to intrinsic features of the parasite genome that complicate assembly, particularly when using short-read sequencing technologies. These include extensive repetitive sequences, a high abundance of transposable elements, large multigene families such as mucins and trans-sialidases,^36^ and substantial variability in genome size and chromosome number across strains.^37^ In addition, frequent aneuploidy and polyploidy, limited synteny, and hybridization events result in highly heterozygous genomes, with early short-read assemblies often failing to fully capture the multiplicity of gene copies present.^38^ Consequently, analyses based on a single genome assembly may overlook or misrepresent members of expanded serine hydrolase families.

To maximize genome coverage, analyses were performed against both the short-read 2014 Dm28c genome^38^ (**Supplementary Data 2**) and the improved long-read-based Dm28c genome assembly reported by Berná *et al.*^26^ (**Supplementary Data 3**). The combined analysis yielded a final set of 37 non-redundant enriched SH-like proteins. These comprised 20 predicted esterases (lipases, phospholipases, deacetylases, acyltransferases), 10 serine peptidases, 1 amidase, and 6 hydrolases of unknown function (**Table 2**). All enriched proteins were supported by *in silico* annotation analysis.

The long-read assembly additionally revealed multiple copies of several enzymes. This is consistent with the known expansion of multigene families in *T. cruzi* such as trans-sialidases, mucins, and mucin-associated surface proteins (MASPs).^26,39,40^ For example, lipase C4B63_104g97 showed the highest copy number with 17 copies, followed by carboxypeptidase CPB1 (C4B63_168g38) with 12 copies. Annotation differences were also observed between assembly versions; for instance, prolyl endopeptidase BCY84_07469 is represented as a full-length sequence in the 2017 assembly but is split into multiple fragments in the 2018 version.

**Table 2.**
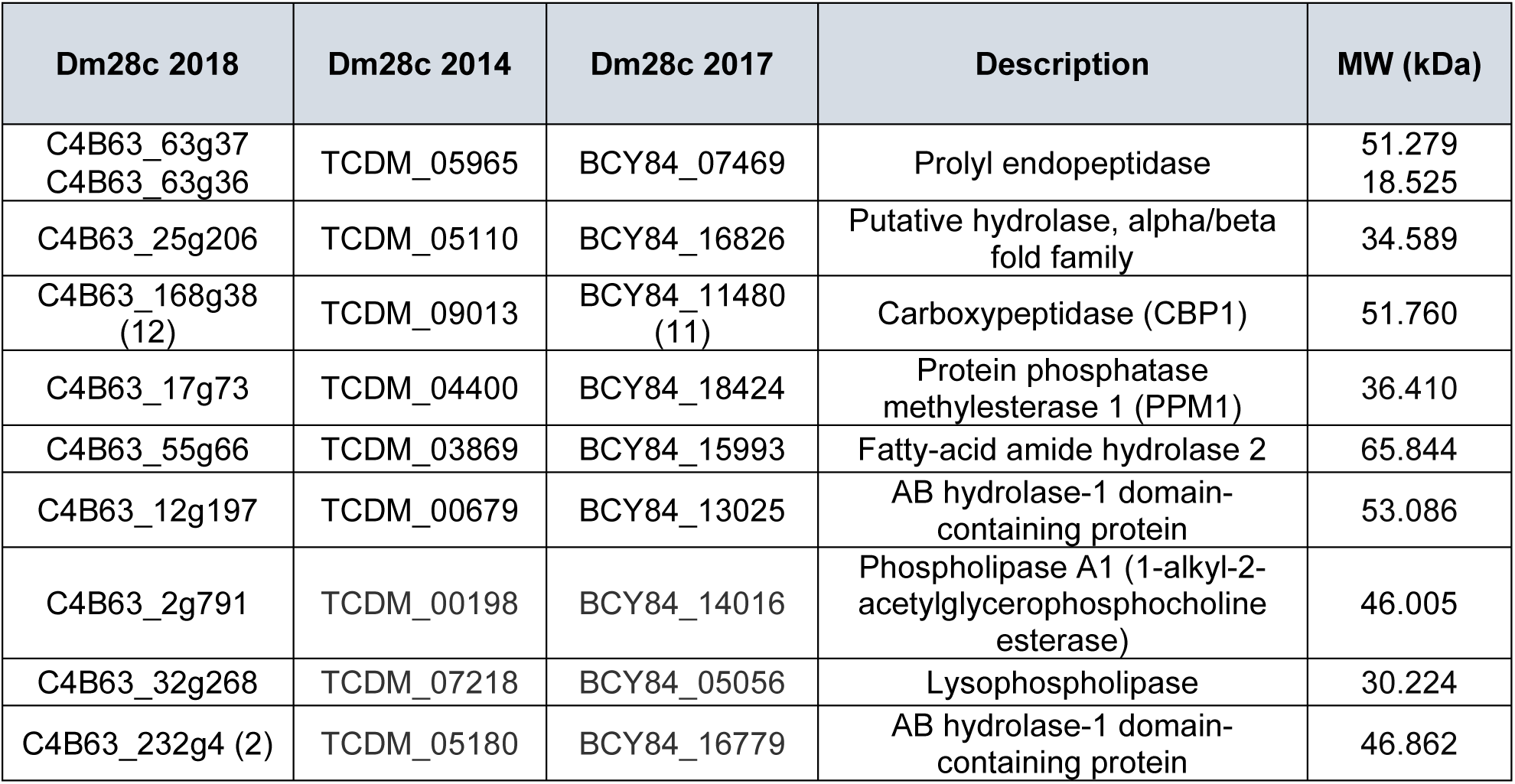
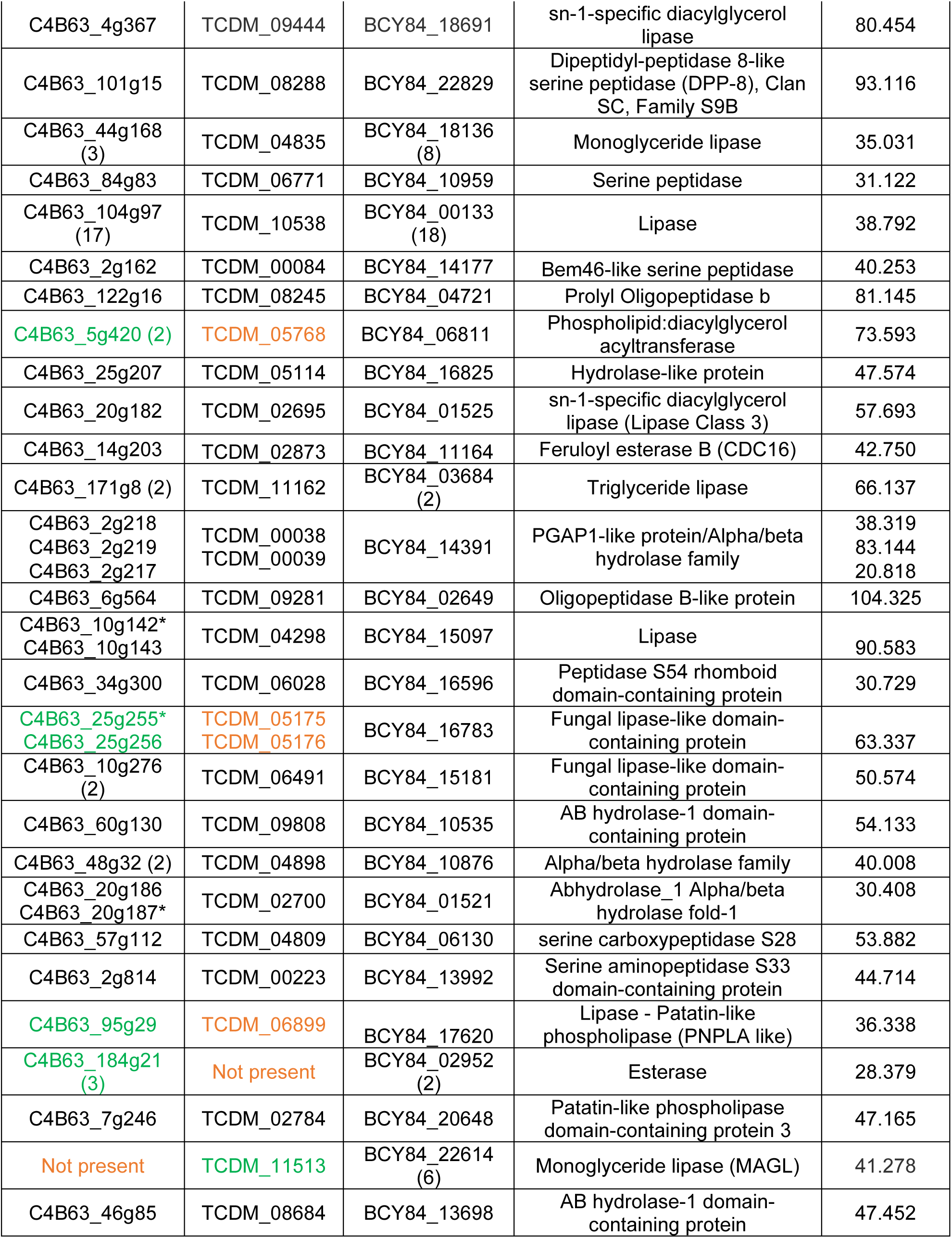
Serine hydrolases identified in *T. cruzi*. Using Dm28c 2018 and 2014 genome assemblies, 35 and 33 SHs were identified respectively, for a total of 37 non-redundant proteins. Proteins identified in only 1 assembly are depicted in green and proteins not enriched or absent from a given assembly are shown in orange The number of copies is shown in parentheses after the accession number, if the copy number is higher than one. Multiple accession numbers per entry indicate that the same gene has been annotated as fragments. *These accession numbers encode part of the gene but they were not detected by chemoproteomics assay.

### Probe selectivity and serinome coverage

Having established the ABPP-enriched serinome, we next evaluated the performance of the FP-alkyne probe panel. With the exception of alkyl probe **1**, which enriched only 3 SHs, all probes displayed broad labelling profiles, enriching more than ten SHs under the conditions tested. Consistently high log_2_ fold changes (≈ 1-7.5; mean ≈ 3.6) together with elevated -log_10_ p-values indicate efficient probe labelling (**Fig. 3a–b**). Probe **7** enriched the highest number of SHs, with 33 targets detected. Probe specificity, defined as the proportion of enriched SHs relative to the total enriched proteins per condition, ranged from 10 % (probe **1**) to 43 % (probe **7**), indicating variable but substantial SH enrichment across the probe series (**Fig. 3** and **Supplementary Fig. S2**).

To determine whether probe enrichment was biased toward highly abundant proteins, intensity-based absolute quantification (iBAQ) values of enriched proteins were compared with genome-wide abundance distributions in *T. cruzi*. Enriched proteins spanned a broad abundance range, including highly abundant enzymes such as serine carboxypeptidase CPB1 (C4B63_168g38 / TCDM_09013) as well as lower-abundance proteins such as prolyl oligopeptidase (C4B63_6g564 / TCDM_09281). These results suggest that probe enrichment is not restricted to the most abundant proteins and support the ability of FP-alkyne probes to capture SHs across a wide dynamic range of protein expression (**Fig. 3b**).

**Figure 3.**
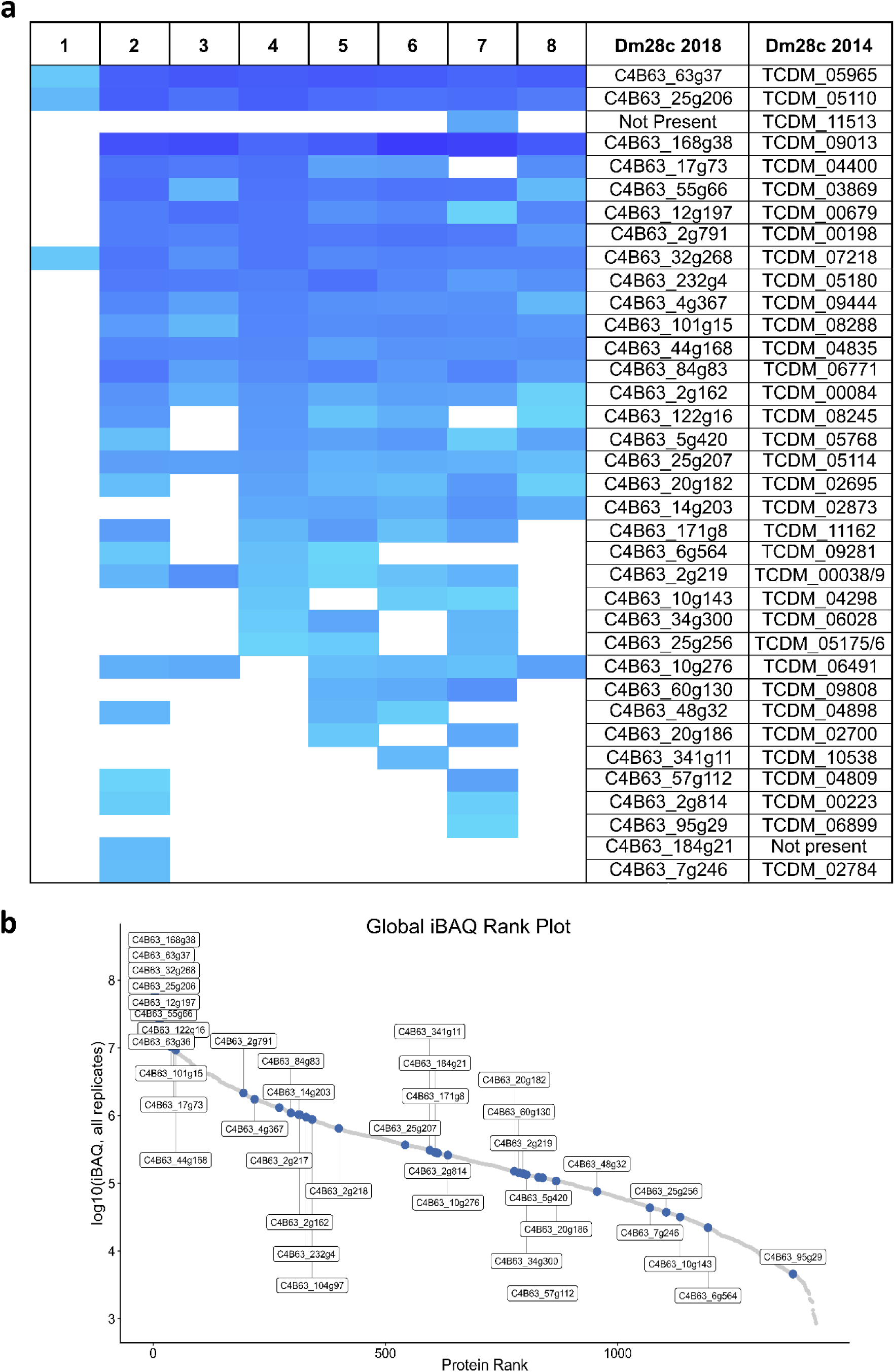
**a.** Log_2_ fold-change (Log_2_ FC) heatmap showing the enrichment profiles of SHs across the different synthesized probes. Log_2_ FC values, reflecting the relative protein abundance between probe-treated and untreated samples, range from 1.2 (light blue) to 7.7 (blue). In brief, *T. cruzi* cultures were treated with the indicated FP-alkyne ABPs, lysed, and subjected to click conjugation with biotin-N_3_. Enriched proteins were subsequently analysed by LFQ LC-MS/MS. **b.** iBAQ abundance distribution of the *T. cruzi* proteome: iBAQ values for all detected proteins were ranked and plotted as log_10_-transformed abundances. SHS enriched by the activity-based probes are highlighted within the overall proteome distribution.

### Pfam domain architecture analysis reveals a diverse repertoire of serine hydrolases in *T. cruzi*

To characterize the enriched SHs, we performed a systematic analysis of predicted protein domain architectures using Pfam annotations, MEROPS classification, conservation across trypanosomatids, and orthologue-based localization data from TrypTagDB (**Fig. 4** and **Supplementary Data S4**). Among the serine peptidase families, member of the S9 family contained either Peptidase_S9_N or DPPIV_N N-terminal domains paired with a C-terminal Peptidase_S9 catalytic domain, consistent with prolyl oligopeptidases (POP) and dipeptidyl peptidases IV (DPPIV). Orthologue-based localization data suggest predominantly cytoplasmic localization for C4B63_63g37, C4B63_122g16, and C4B63_101g15, whereas C4B63_6g564 showed partial mitochondrial localization (50%), suggesting the possibility of dual targeting. The S10 family member C4B63_168g38, annotated as a serine carboxypeptidase, showed a striking expansion, with 12 predicted gene copies sharing a conserved domain architecture, consistent with the broader multi-gene family amplification observed in *T. cruzi* for trans-sialidases, mucins, and MASPs.^26,39,40^

**Figure 4.**
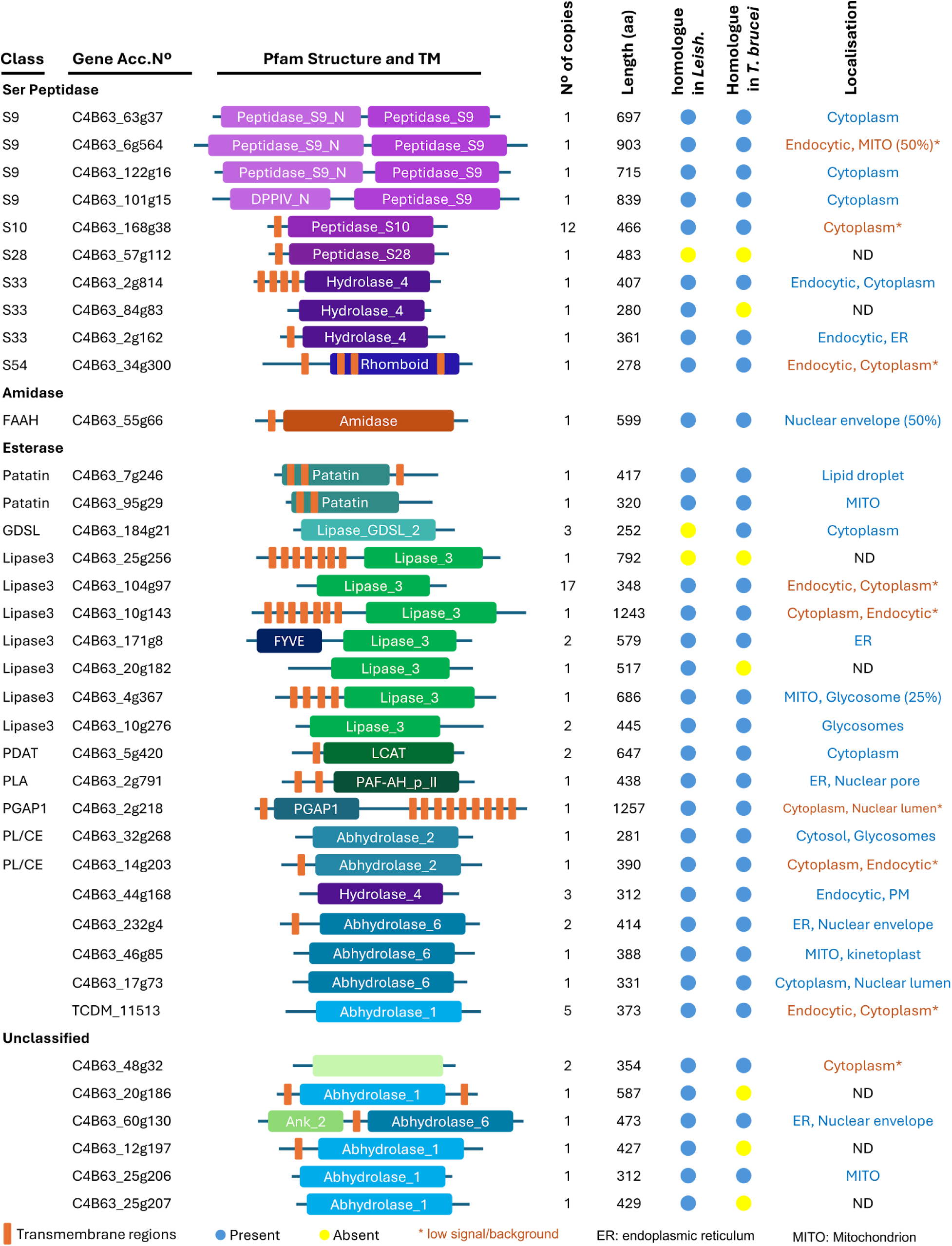
Pfam analysis of enriched SHs. The figures are not drawn to scale; only the relative positions of the domains and TM regions are shown. The data for this figure are presented in Supplementary Data S4. Localization was obtained from experimental data in TrypTagDB.^42^ FAAH: Fatty-acid amide hydrolase, Patatin: Patatin-like Lipase, GDSL: GDSL Lipase, Lipase3: Lipase class 3, PDAT: Phospholipid:diacylglycerol acyltransferase, PLA: Phospholipase A, PGAP1: GPI inositol-deacylase PGAP1, PL/CE: Phospholipase/Carboxylesterase, ER: endoplasmic reticulum, MITO: mitochondrion, and PM: plasma membrane.

### Gene Ontology enrichment and protein interaction network analysis support lipid metabolism as a dominant theme in the ABPP-enriched serinome

To further assess the chemoproteomics dataset and explore the collective biological relevance of the ABPP-enriched SHs, we performed Gene Ontology (GO) enrichment and protein-protein interaction (PPI) network analyses using the STRING database, applying the *T. cruzi* CL Brener proteome as the reference background. The CL Brener strain proteome was used because it is the only *T. cruzi* assembly available in STRING; enriched Dm28c proteins were therefore mapped to their CL Brener homologues for the GO and PPI analyses.

GO enrichment revealed that the most significantly enriched molecular functions included “hydrolase activity,”, “carboxylic ester hydrolase activity”, and “serine-type peptidase activity”, supporting the high selectivity of the ABPs toward SHs (**Fig. 5a** and **Supplementary Data 5**). Among biological processes, “lipid metabolic process”, “lipid catabolic process”, and “acylglycerol catabolic process” were strongly overrepresented, indicating that many of the enriched enzymes are associated with lipid turnover and energy mobilization. This is consistent with structural diversity of the lipase families captured and with the prevalence of predicted transmembrane regions and inferred organelle-associated architectures described above. Cellular component annotations further indicated that a substantial subset of targets is associated with membranes, supporting the prevalence of membrane-embedded and organelle-associated SHs.

PPI analysis revealed a tightly interconnected network among the enriched proteins (**Fig. 5b**), with strong associations involving nodes annotated as monoglyceride lipase, lysophospholipase, and prolyl oligopeptidase. These enzymes are, again, associated with lipid metabolism, membrane remodelling, and peptide processing. The overall network exhibited highly significant enrichment (PPI enrichment p-value < 1 × 10^-16^), indicating that the identified proteins are biologically connected rather than representing a random sampling of the proteome. Together, the GO and PPI analyses underscore the functional coherence of the captured dataset and indicate that lipid metabolism is a dominant biological theme within the active SH landscape of *T. cruzi* epimastigotes.

These functional themes are consistent with the known lipid auxotrophy of trypanosomatids. Unable to synthesize certain lipids *de novo*, *T. cruzi* and related parasites rely heavily on host lipid acquisition and remodelling, with lipid metabolism linked to parasite differentiation, intracellular survival, and virulence.^41,42^ The enrichment of phospholipase A, lysophospholipase, patatin-like phospholipases, and multiple lipase 3 family members suggests that SHs may contribute broadly to these lipid-handling processes. The inferred glycosomal or mitochondrial association of multiple lipases is also noteworthy, as glycosomes are vital trypanosomatid-specific organelles that compartmentalize key metabolic pathways, including glycolysis and aspects of lipid metabolism.^43–46^

**Figure 5.**
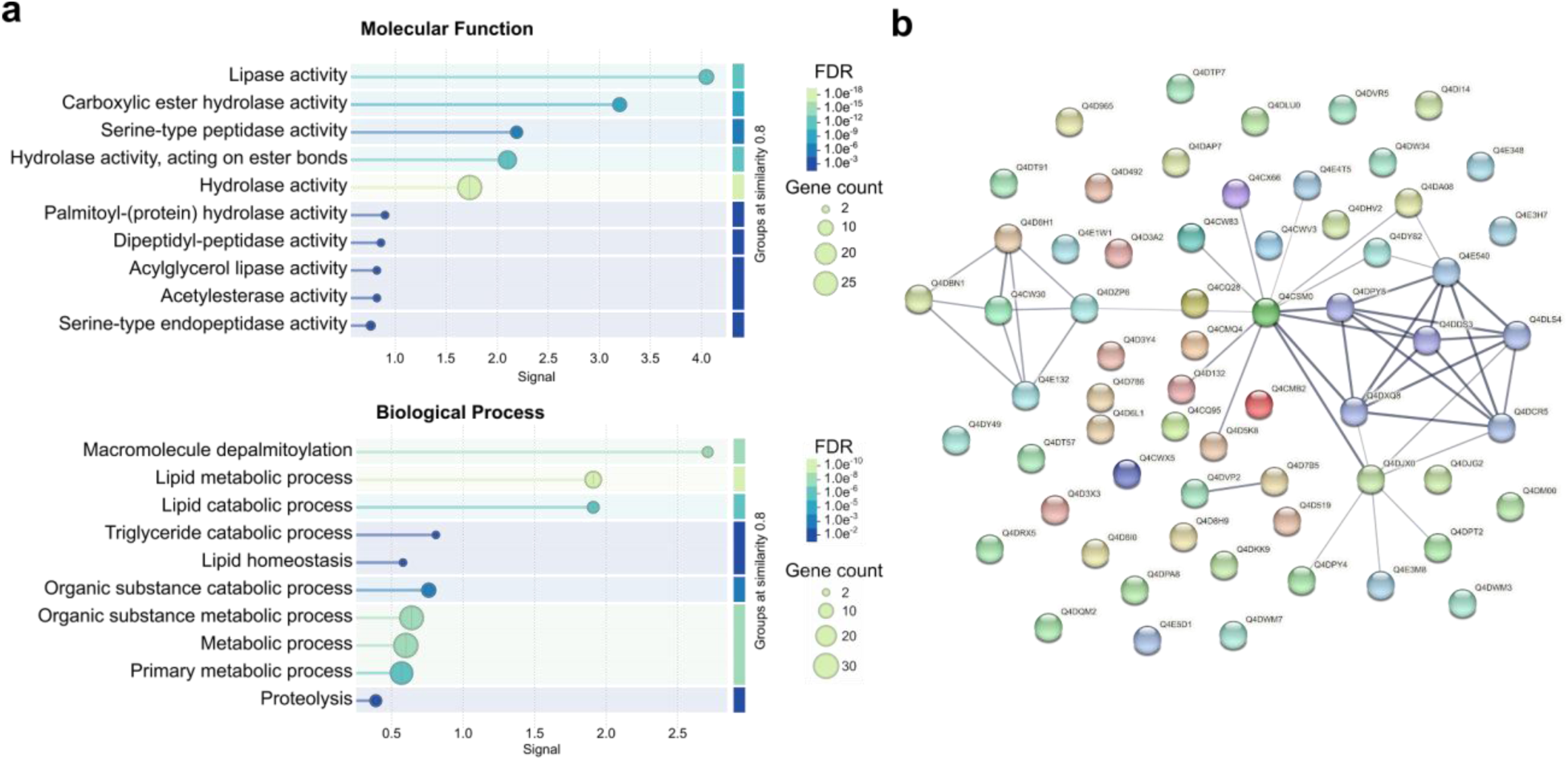
**a.** Gene ontology (GO) analysis for the proteins enriched by the ABPs. The enrichment analysis was performed using STRING database v12.0^47^, applying a one-sided Fisher’s exact test against the organism-specific background proteome. Resulting p-values were corrected for multiple testing using the Benjamini–Hochberg FDR, and enriched terms were ranked according to their FDR values. **b.** Protein– protein interaction (PPI) network of enriched SHs generated with STRING database v11.5^47^

## Discussion

Combining live-cell ABPP with genome-informed curation, we generated the first activity-based map of the *T. cruzi* epimastigote serinome, resolving 37 enriched SH-like proteins, of which 35 retain a conserved or partially conserved catalytic triad or dyad. Whole-cell labelling with cell-permeable FP-alkyne probes was the key enabling step: lysate-based labelling performed poorly, consistent with extract agglutination and restricted probe access, whereas profiling intact parasites both circumvented this barrier and captured enzymes in their native activity state, minimizing the artifactual activation or inactivation of labile hydrolases that lysis can introduce. Together, these results illustrate how live-cell FP-ABPP can be applied to kinetoplastid parasites whose dense surface architecture frustrates conventional lysate-based labelling, and suggest this whole-cell strategy may be broadly useful for chemoproteomic profiling in such systems. The 35 active SHs recovered correspond to ca. 63 % of the 56 SHs predicted *in silico*, and represents substantial coverage for an activity-based experiment, and consistent with FP probes reporting selectively on the active, non-zymogen fraction of the family. The identity of the uncaptured 37 % is biologically informative in itself. Several of these enzymes might be absent in the epimastigote stage, which is the insect-adapted proliferative form and is metabolically and proteomically distinct from the clinically relevant trypomastigote and intracellular amastigote stages. This possibility is consistent with our previous activity-based profiling studies in Leishmania, where enzyme activity landscapes varied across life cycle contexts.^17,29^ Others may exist as catalytically inactive zymogens awaiting specific activating signals, or may target substrates not accessible to FP-based probes, such as enzymes with occluded active sites. This interpretation reframes the uncaptured fraction not as a limitation of the approach, but as a biological signal highlighting the depth of SH regulation across the *T. cruzi* life cycle. Future ABPP profiling of trypomastigotes and amastigotes, particularly the disease-critical intracellular amastigote, would be expected to reveal additional active SHs and provide a stage-resolved view of serinome dynamics.

Although the biochemical roles of many of the SHs detected remain to be established, several identified enzymes have well-characterized functions in parasite virulence or host-pathogen interactions. Among the serine peptidases, three prolyl endopeptidases, C4B63_63g36-7 (TCDM_05965), C4B63_122g16 (TCDM_08245), and C4B63_6g564 (TCDM_09281), have been previously characterized as virulence factors in both *T. cruzi* and *Leishmania* spp., with orthologs implicated in parasite-host processes.^9,18–20,48–51^ More specifically, C4B63_122g16 (TCDM_08245), known as oligopeptidase B (OPB), belongs to the prolyl oligopeptidase family and has been associated with trypomastigote penetration into mammalian cells.^9,52,53^ The carboxypeptidase CPB1 (C4B63_168g38/TCDM_09013), also known as *Tc*SCP, is a lysosomal glycoprotein serine carboxypeptidase identified as a minor antigen recognized by sera from patients with chronic Chagas disease.^9,54,55^ Together with Tc80, these enzymes have been proposed or investigated as candidate therapeutic targets in Chagas disease,^48^ and their enrichment here supports the ability of the workflow to recover biologically relevant SHs. Notably, these enzymes were detected as catalytically active in the epimastigote stage profiled here, whereas their characterized roles in mammalian-cell invasion and host interaction have been established in infective *T. cruzi* stages or in orthologs from related trypanosomatids.

The domain architecture analysis also provided insight into the cellular context in which these enzymes may act. The prevalence of predicted integral membrane topologies, particularly among lipase 3, S33 peptidase, and PGAP1-like family members, suggest that a substantial fraction of the *T. cruzi* serinome may act at membrane interfaces, potentially processing lipidic or membrane-anchored substrates. Notably, a substantial proportion of the enriched SHs are associated with lipid metabolism (54%). This is significant as trypanosomatids, including *T. cruzi*, are unable to synthesize certain lipids and therefore rely on lipid uptake from their hosts and consequently these lipases may represent potential targets for therapeutic intervention.^41,42^ For example, lysophospholipase C4B63_32g268 (TCDM_07218) has been implicated in virulence and pathogenicity across diverse organisms, including bacteria and fungi.^56,57^ This enzyme has homologs identified in *T. brucei*^59^ and *L. mexicana*. Inhibition of a related enzyme class has also been shown to block *Plasmodium falciparum* replication *in vitro*.^58^ Among the other lipases identified here, a phospholipase A1 (C4B63_2g791/TCDM_00198; also known as PLA1) was enriched. Phospholipases A1 have been described as virulence-associated enzymes that participate in the early stages of parasite–host interactions preceding invasion.^56,57,59–61^ Their activity can alter the host-cell lipid profile by generating second messengers, including diacylglycerol, free fatty acids, and lysophosphatidylcholine, which activate the protein kinase C signalling pathway and promote parasite invasion.^59–61^ However, the phospholipase A1 identified in this study does not appear to correspond directly to the phospholipase homologues previously characterized in *Trypanosoma* spp suggesting that it may represent a distinct family member with a potentially different biological role.

Two findings are particularly noteworthy. Firstly, C4B63_171g8 combines a FYVE zinc finger domain with a class 3 lipase (Lipase_3) catalytic domain, suggesting recruitment to PI3P-enriched endosomal membranes and spatial regulation of hydrolytic activity. This architecture is uncommon but it has been previously predicted in *Arabidopsis thaliana*.^62,63^ To our knowledge, this combination of domains has not been previously reported in trypanosomatids. Second, the enrichment of a catalytically supported rhomboid intramembrane serine protease (C4B63_34g300/TCDM_06028) is notable given the essential role of rhomboid proteases in invasion in apicomplexan parasites.^64–66^ These architectural features underscore that the *T. cruzi* serinome includes not only canonical soluble hydrolases but also membrane-associated and mechanistically unusual enzymes that warrant further functional and chemical-biology investigation.

Finally, we examined conservation of the enriched proteins across the related trypanosomatids *T. brucei* and *Leishmania*. Of the 37 enriched proteins, 29 were present in both related trypanosomatid lineages, one was absent from *Leishmania,* and five were absent in *T. brucei*. Two proteins, S28 peptidase C4B63_57g112 and the lipase 3 enzyme C4B63_25g256, were absent from both *T. brucei* and *Leishmania* (**Fig. 4**), suggesting that they may represent candidates for studies of *T. cruzi*-restricted SH function.^a^

To place these observations in a broader evolutionary context, we analysed co-occurrence of the identified SHs across representative genomes from the three domains of life (Eukaryota, Archaea, and Bacteria) using STRING database. As shown in **Supplementary Figure S3**, most enriched SHs were broadly conserved across trypanosomatid genomes and eukaryotes more generally, with reduced representation in bacteria and near-absence in archaea. This pattern is consistent with the eukaryotic origin of most SH families identified here. From a target prioritization perspective, SHs that are conserved across trypanosomatids but divergent from human orthologs, for example oligopeptidase b, may be of particular interest.

## Conclusions

This study further reinforces the role ABPP has in early-stage drug discovery. This study provides the first activity-based chemoproteomic functional map of the *T. cruzi* epimastigote serinome, identifying 35 enriched SH-like proteins, with conserved catalytic triad/dyad features, through a live-cell ABPP-LFQ workflow. The dataset highlights lipid-metabolism-associated hydrolases, virulence-associated peptidases, and previously uncharacterized enzymes as candidates for future functional validation. By integrating active-site-directed chemical probes with genome-informed annotation, this work establishes a reference resource for stage-resolved *T. cruzi* serinome biology. Beyond its value as a biological annotation resource, this dataset has direct relevance for chemical biology and antiparasitic target discovery. Serine hydrolases are among the most chemically tractable enzyme classes, and FP-based probes report active-site accessibility in a cellular context. In alignment with the WHO NTD 2021–2030 roadmap objectives for new molecular targets in Chagas disease,^7^ the enriched enzymes identified here therefore represent a prioritized set of probe-accessible and catalytically competent targets for future inhibitor screening,^67^ competitive ABPP, and structure-guided ligand discovery. Questions remain as to the role of SH in trypomastigotes and intracellular amastigotes and the specific function, essentiality, and therapeutic relevance of the identified enzymes. Work addressing these important questions is currently underway and will be reported in due course.

## Acknowledgements

We thank Dr. Germán L. Rosano and the IBR-CONICET MS facility for the MS analysis. We also thank Dr. Max Ruwolt for his input and assistance with the development of the R scripts.

## Author Contributions

Conceptualization, Jaime A. Isern, Exequiel O.J. Porta, Julia A. Cricco, Guillermo R. Labadie, and Patrick G. Steel; Methodology, Jaime A. Isern, Exequiel O.J. Porta; Formal Analysis, Jaime A. Isern, Marcelo L. Merli, and Exequiel O.J. Porta; Investigation, Jaime A. Isern, Maria Gabriela Mediavilla, Maria Sol Ballari; Writing, Original Draft Preparation, Jaime A. Isern, Marcelo L. Merli, and Patrick G. Steel; Funding Acquisition, Patrick G. Steel, Guillermo R. Labadie, and Julia A. Cricco; Writing, Review & Editing, all authors. All authors have read and agreed to the published version of the manuscript.

## Data Availability Statement

The data that support the findings of this study are available in the supplementary material of this article.

The mass spectrometry proteomics data have been deposited to the ProteomeXchange Consortium via the PRIDE^68^ partner repository with the dataset identifier PXD080813.

## Code Availability Statement

The R scripts used in this study are available without restrictions via Zenodo: https://doi.org/10.5281/zenodo.20626073

R scripts were adapted by the authors from open source scripts.^69^ ChatGPT (OpenAI) was used to assist with code optimization and troubleshooting. All final scripts were manually reviewed, tested, and validated by the authors.

## Footnotes

a They may be restricted to *T. cruzi* and closely related species such as *Trypanosoma theileri*, as well as the ancestral free-living non-parasitic *Bodo saltans*.

