## Supplementary Information for "Functional Mapping of the *Trypanosoma cruzi* Serinome by Fluorophosphonate Activity-Based Protein Profiling"

#### SUPPLEMENTARY DATA

**Supplementary Data 1:** Details of the Serine Hydrolases searches *in silico* in *T. cruzi* strain Dm28c 2018 genome and Manual curation and classification of the identified genes.

**Supplementary Data 2:** Proteomic results obtained using Dm28c 2014 genome

**Supplementary Data 3:** Proteomic results obtained using Dm28c 2018 genome

**Supplementary Data 4**: Details of the information shown in Fig. 3. The table contains the Pfam of each enriched protein, the accession numbers of the *Leishmania* and *T. brucei* homologues, and localization according to TrypTagDB.

**Supplementary Data 5**: Results of the GO analyses.

### SUPPLEMENTARY FIGURES

#### Supplementary Figure S1

**PeptidaseS28 (C4B63_17g176)**

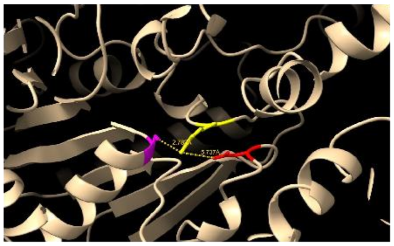

S179 R438 D411

Complete sequence and structure taken from accession number TcCLB.506425.10. Interatomic distances 2.783 and 5.737 A (partially conserved). The pLDDT are 94.44, 88.19, and 92.88, respectively.
Model: AF-Q4DM56-F1-v6

**PeptidaseS28 (C4B63_57g112)**

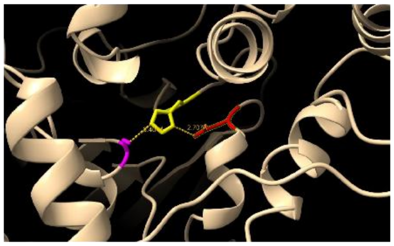

S193 H451 D424

Interatomic distances 3.404 and 2.707 Å. The pLDDT are 96.81, 94.31, and 95.69, respectively.
Model: AF-A0A2V2V000-F1-v6

**Peptidase S33 (C4B63_2g814)**

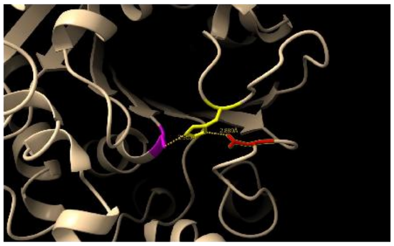

S271 H385 D347

Interatomic distances 2.953 and 2.889 Å. The pLDDT are 97.94, 96.25, and 96.38, respectively.
Model: AF-A0A2V2W413-F1-v6

**Peptidase S33 (C4B63_84g83)**

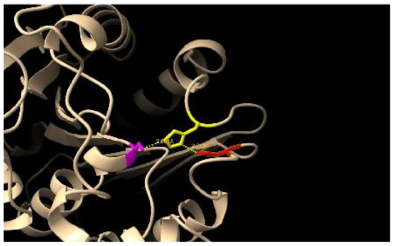

S138 H245 D207

Interatomic distances 2.861 and 2.719 Å. The pLDDT are 95.69, 94.19, and 95.50, respectively.
Model: AF-A0A2V2UU76-F1-v6

**PeptidaseS33 (C4B63_2g162)**

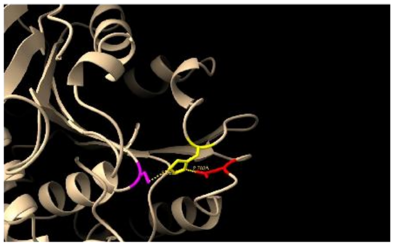

S209 H333 D301

Interatomic distances 3.052 and 2.783 Å. The pLDDT are 97.50, 96.31, and 97.19, respectively.
Model: AF-A0A2V2W2I3-F1-v6

**PeptidaseS51 (C4B63_27g229)**

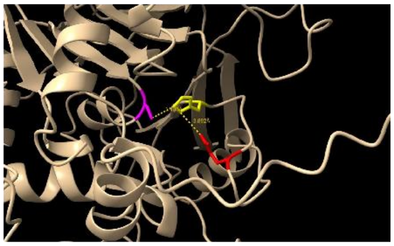

S144 H228 E233

Interatomic distances 3.105 and 3.892 Å. The pLDDT are 98.25, 97.88, and 89.19, respectively.
Model: AF-A0A2V2VFP5-F1-model_v6

**Peptidase S54 (C4B63_34g300)**

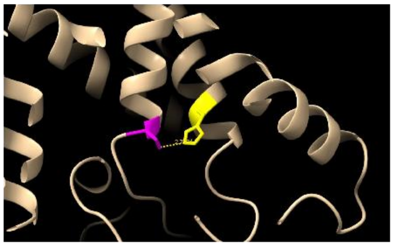

S185 H244

Interatomic distance 2.732 Å. The pLDDT are 91.75 and 91.00, respectively.
Model: AF-A0A2V2V9H6-F1-v6

**Fatty-acid amide hydrolase (FAAH) (C4B63_55g66)**

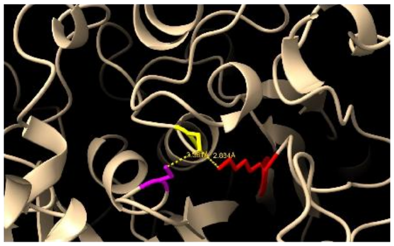

S244 S220 K145

Interatomic distances 3.381 and 2.884 Å. The pLDDT are 98.06, 97.81, and 98.75, respectively.
Model: AF-A0A2V2V0U3-F1-v6

**Patatin lipase (C4B63_18g341)**

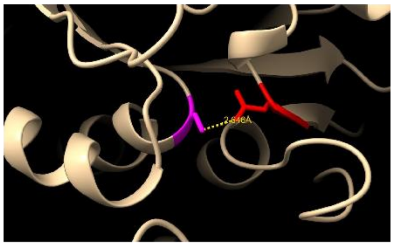

S370 D523

Interatomic distance 2.646 Å. The pLDDT are 90.56 and 84.25, respectively.
Model: AF-A0A2V2VJI4-F1-v6

**Patatin lipase (C4B63_7g246)**

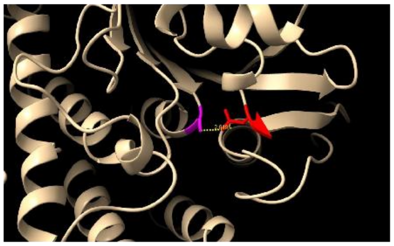

S58 D184

Interatomic distance 2.649 Å. The pLDDT are 94.81 and 91.25, respectively.
Model: AF-A0A2V2W0N3-F1-v6

**GDSL Lipase (C4B63_184g21)**

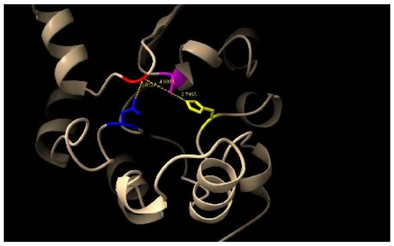

N83 G46 S15 H206

Interatomic distances 3.852, 4.593, and 2.790 Å. The pLDDT are 98.25, 97.25, 98.62 and 98.50, respectively.
Model: AF-A0A2V2ULF1-F1-v6

**GDSL Lipase (C4B63_13g128)**

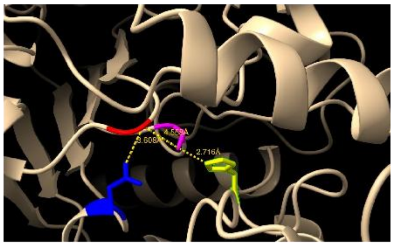

N367 G334 S251 H519

Interatomic distances 3.608, 4.558, and 2.716 Å.The pLDDT are 94.00, 84.00, 92.81, and 96.19, respectively.Model: AF-A0A2V2VPA7-F1-v6

**Peptidase S1 (C4B63_23g231)**

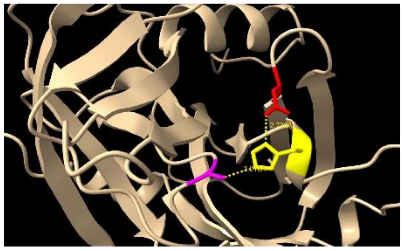

S149 H49 D83

Interatomic distances 3.018 and 2.738 Å. The pLDDT are 91.38, 92.12, and 94.31, respectively. Model: AF-A0A2V2VIB2-F1-v6

**PeptidaseS1 (C4B63_9g401)**

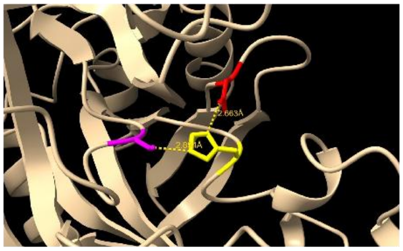

S236 H57 D114

Interatomic distances 2.894 and 2.663 Å.The pLDDT are 96.00, 94.81, and 96.38, respectively. Model: AF-A0A2V2VVB8-F1-v6

**PeptidaseS26 (C4B63_31g105)**

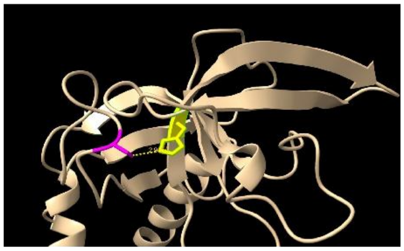

S86 H124

Interatomic distances 2.955 Å.The pLDDT are 92.00, and 96.31, respectively. Model: AF-A0A2V2VD34-F1-v6

**PeptidaseS26 (C4B63_13g150)**

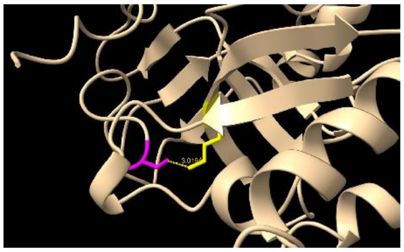

S35 K84

Interatomic distance 3.019 Å.The pLDDT are 94.19 and 96.75, respectively. Model: AF-A0A2V2VP91-F1-v6

**Peptidase S8 (C4B63_10g296)**

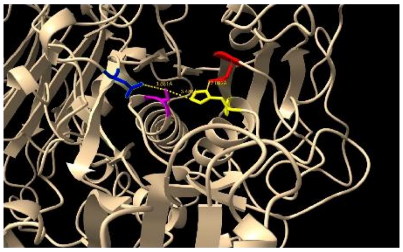

N531 S612 H430 D378

In the absence of an available structure for this protein, the structure corresponding to accession number TcCLB.509669.20 was used as a representative model. Interatomic distances 3.861, 3.486, and 2.892 Å. The pLDDT are 83.00, 95.06, 92.12, and 93.69, respectively. Model: AF-Q4DP21-F1-v6

**PeptidaseS8 (C4B63_95g13)**

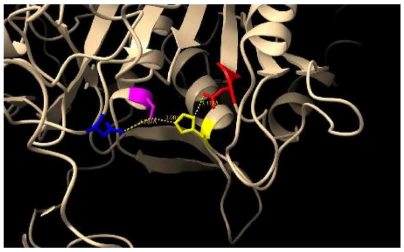

N370 S480 H246 D196

In the absence of an available structure for this protein, the structure corresponding to accession number TcCLB.511859.60. Interatomic distances 4.737, 3.001, and 3.178 Å. The pLDDT are 86.62, 94.38, 91.62 , and 96.19, respectively. Model: AF-Q4DLQ6-F1-v6

**PeptidaseS9 (C4B63_63g37)**

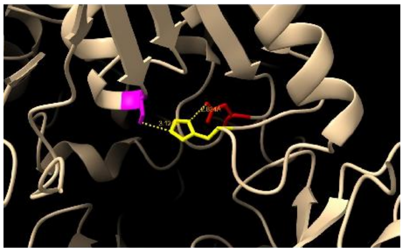

S548 H667 D631

Complete sequence and structure taken from accession number TcCLB.506247.230. Interatomic distances 3.121 and 2.834 Å. The pLDDT are 98.44, 88.25, and 94.94, respectively. Model: AF-Q4E132-F1-v6

**PeptidaseS9 (C4B63_6g564)**

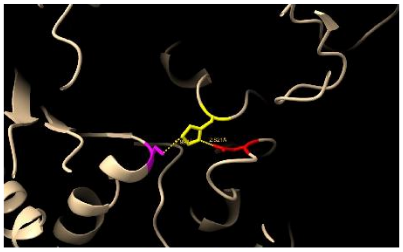

S629 H758 D717

Interatomic distances 2.997 and 2.821 Å.The pLDDT are 96.00, 84.75, and 87.50, respectively. Model: AF-A0A2V2VWU5-F1-v6

**PeptidaseS9 (C4B63_122g16)**

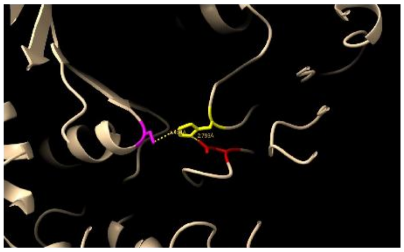

S563 H683 D648

Interatomic distances 3.494 and 2.793 Å.The pLDDT are 97.69, 89.62, and 94.62, respectively. Model: AF-A0A2V2UT74-F1-v6

**PeptidaseS9 (C4B63_101g15)**

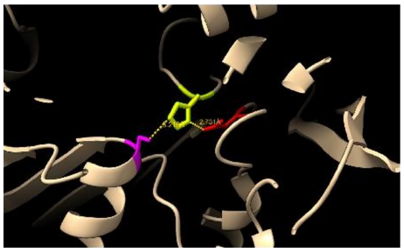

S692 H815 D783

Interatomic distances 3.218 and 2.731 Å.The pLDDT are 97.19, 94.12, and 94.00, respectively.Model: AF-A0A2V2USR9-F1-v6

**PeptidaseS10 (C4B63_168g38)**

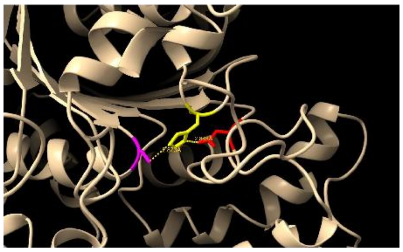

S182 H442 D379

Interatomic distances 2.873 and 2.843 Å.The pLDDT are 98.62, 98.81, and 98.88, respectively. Model: AF-A0A1B1R0W4-F1-v6

**PeptidaseS15 (C4B63_6g235)**

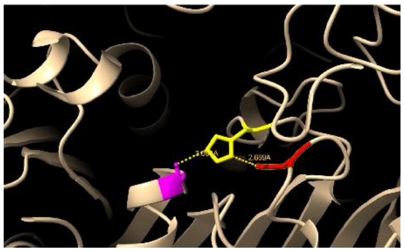

S125 H272 D246

Interatomic distances 3.093 and 2.669 Å.The pLDDT are 96.69, 96.50, and 95.88, respectively.Model: AF-A0A2V2VWK6-F1-v6

**Lipase 3 (C4B63_25g256)**

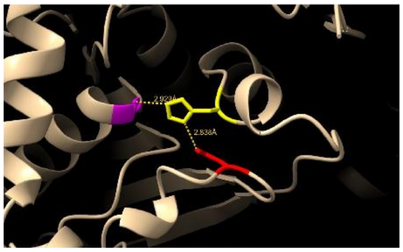

S685 H762 D736

Complete sequence and structure taken from accession number TcCLB.507047.90. Interatomic distances 2.929 and 2.838 Å. The pLDDT are 88.94, 85.50, and 80.06, respectively.
Model: AF-Q4DPS8-F1-v6

**Lipase class 3 (C4B63_22g182)**

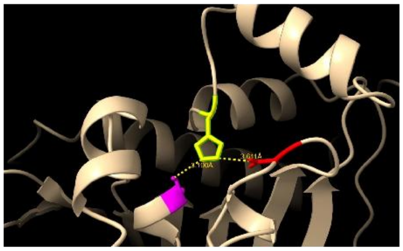

S375 H492 D450

Interatomic distances 3.190 and 3.611 Å. The pLDDT are 86.44, 63.22, and 69.44, respectively.
Model: AF-A0A2V2UKM7-F1-model_v6

**Lipase 3 (C4B63_104g97)**

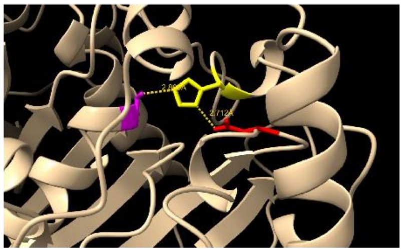

S157 H284 D221

Interatomic distances 2.865 and 2.712 Å. The pLDDT are 96.62, 93.56, and 94.94, respectively.
Model: AF-A0A2V2UVW5-F1-v6

**Lipase 3 (C4B63_10g143)**

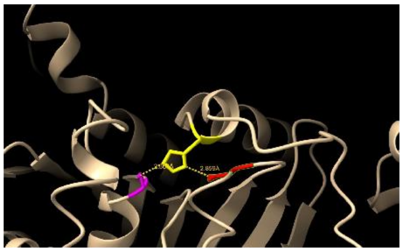

S1078 H1188 D1139

Complete sequence and structure taken from accession number TcCLB.509999.50. Interatomic distances 2.864 and 2.859 Å. The pLDDT are 87.62, 84.94, and 78.81, respectively.
Model: AF-Q4DKK9-F1-v6

**Lipase 3 (C4B63_171g8)**

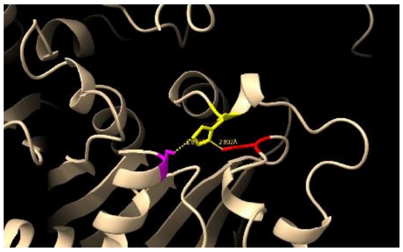

S380 H563 D441

Interatomic distances 2.877 and 2.802 Å. The pLDDT are 89.75, 83.12, and 84.12, respectively.
Model: AF-A0A2V2UMG9-F1-v6

**Lipase 3 (C4B63_10g145)**

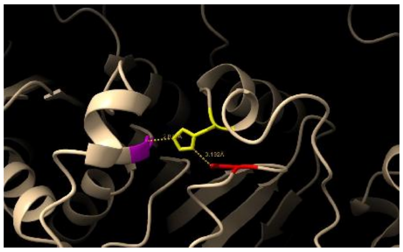

S1118 H1227 D1179

Complete sequence and structure taken from accession number TcCLB.509999.60. Interatomic distances 2.872 and 3.192 Å. The pLDDT are 88.88, 84.00, and 77.12, respectively.
Model: AF-Q4DKK8-F1-v6

**Lipase 3 (C4B63_20g182)**

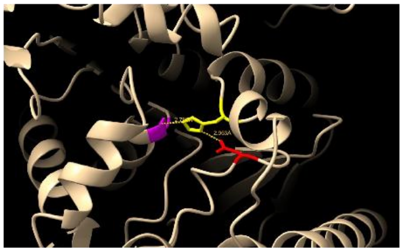

S350 H489 D410

Interatomic distances 2.793 and 2.953 Å. The pLDDT are 91.94, 88.06, and 88.69, respectively.
Model: AF-A0A2V2VHS3-F1-v6

**Lipase 3, C4B63_4g367**

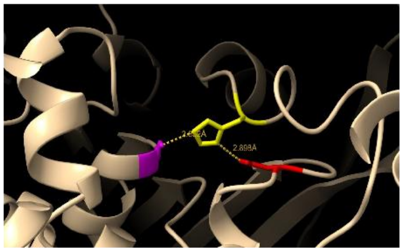

S451 H642 D507

Complete sequence and structure taken from accession number TCDM_09444. Interatomic distances 2.852 and 2.898 Å. The pLDDT are 96.31, 95.50, and 95.44, respectively.
Model: AF-V5APZ1-F1-v6

**Lipase 3 (C4B63_20g60)**

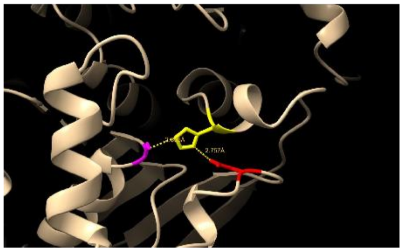

S773 H850 D824

Interatomic distances 2.895 and 2.757 Å. The pLDDT are 89.38, 85.56, and 82.31, respectively.
Model: AF-A0A2V2VNF6-F1-v6

**Lipase 3 (C4B63_10g276)**

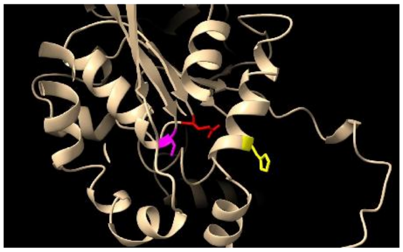

S150 H283 E171

Lacks the functional orientation. The pLDDT are 87.81, 72.88, and 90.00, respectively.
Model: AF-A0A2V2VFX4-F1-v6

**Phospholipid:diacylglycerol acyltransferase (PDAT) (C4B63_5g420)**

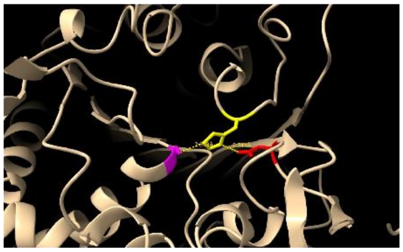

S348 H592 D543

Interatomic distances 2.849 and 2.741 Å. The pLDDT are 97.00, 93.00, and 94.69, respectively.
Model: AF-A0A2V2W3E4-F1-v6

**Phospholipase A (C4B63_2g791)**

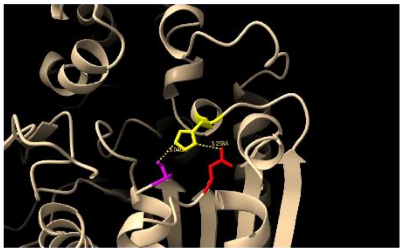

S277 H387 D308

Complete sequence and structure taken from accession number TCDM_00198. Interatomic distances 3.046 and 3.258 Å. The pLDDT are 97.44, 91.00, and 97.19, respectively.
Model: AF-V5BD01-F1-v6

**GPI inositol-deacylase (PGAP1) (C4B63_2g218)**

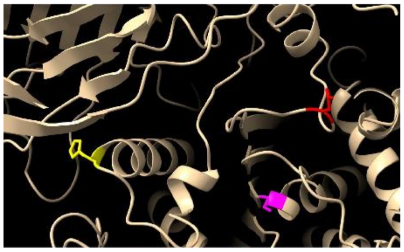

S300 H454 D406

Complete sequence and structure taken from accession number TcCLB.506147.170. Structure lacks the functional orientation.The pLDDT are 63.03, 44.44, and 27.17, respectively.
Model: AF-Q4DT91-F1-v6

**GPI inositol-deacylase (PGAP1) (C4B63_23g274)**

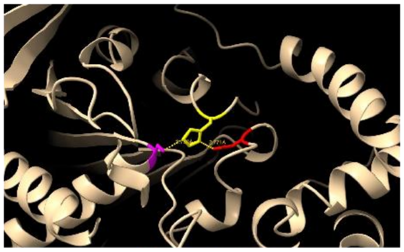

S260 H376 D339

Interatomic distances 2.789 and 2.771 Å. The pLDDT are 93.88, 89.12, and 83.75, respectively.
Model: AF-A0A2V2VI68-F1-v6

**LID Hydrolase (C4B63_8g548)**

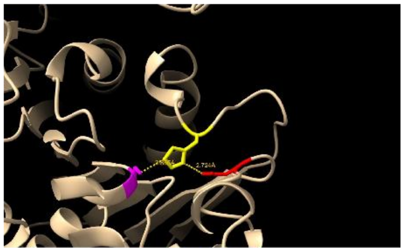

S134 H296 D265

Interatomic distances 2.865 and 2.724 Å. The pLDDT are 95.38, 93.69, and 92.00, respectively.
Model: AF-A0A2V2VTB9-F1-v6

**Phospholipase/Carboxylesterase (C4B63_32g268)**

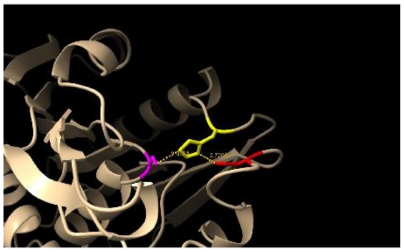

S172 H256 D224

Interatomic distances 2.925 and 2.720 Å.The pLDDT are 97.38, 96.38, and 97.12, respectively.Model: AF-A0A2V2V9U5-F1-v6

**Phospholipase/Carboxylesterase (C4B63_14g203)**

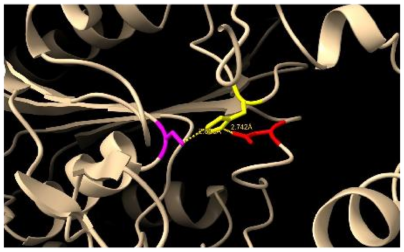

S203 H365 D267

Interatomic distances 2.806 and 2.742 Å. The pLDDT are 97.81, 96.88, and 97.56, respectively.
Model: AF-A0A2V2VU40-F1-v6

**Bifunctionalcarbohydrateesterase (CE) (C4B63_83g76)**

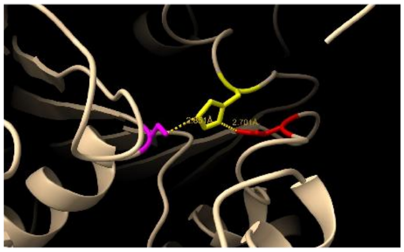

S337 H514 D460

Interatomic distances 2.851 and 2.701 Å. The pLDDT are 93.50, 85.44, and 87.62, respectively.
Model: AF-A0A2V2UX88-F1-v6

**Unclassified Esterase (C4B63_44g168)**

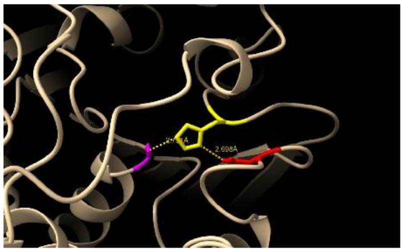

S139 H286 D256

Interatomic distances 2.791 and 2.698 Å. The pLDDT are 97.19, 96.19, and 94.81, respectively.
Model: AF-A0A2V2V407-F1-v6

**Unclassified Esterase (C4B63_232g4)**

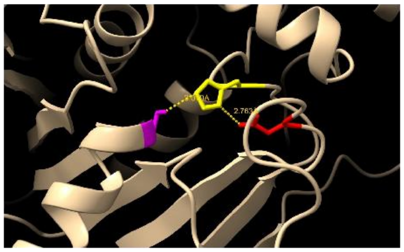

S211 H381 D350

Interatomic distances 3.070 and 2.763 Å. The pLDDT are 97.56, 97.12, and 97.06, respectively.
Model: AF-A0A2V2UMB8-F1-v6

**UnclassifiedEsterase (C4B63_46g85)**

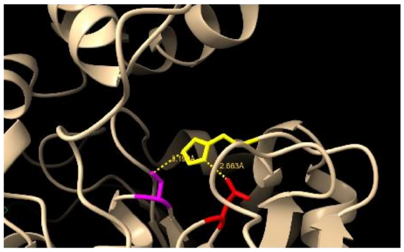

S105 H358 D205

Interatomic distances 3.190 and 2.663 Å. The pLDDT are 94.56, 91.75, and 96.44, respectively.
Model: AF-A0A2V2V610-F1-v6

**Unclassified Esterase (C4B63_17g73)**

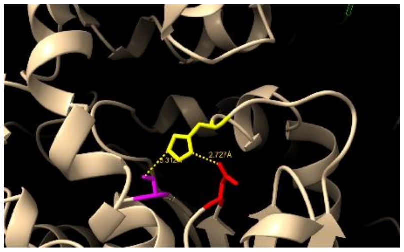

S142 H295 D169

Interatomic distances 3.312 and 2.727 Å.The pLDDT are 95.62, 88.50, and 94.31, respectively.Model: AF-A0A2V2VKK1-F1-v6

**Unclassified Esterase (C4B63_6g260)**

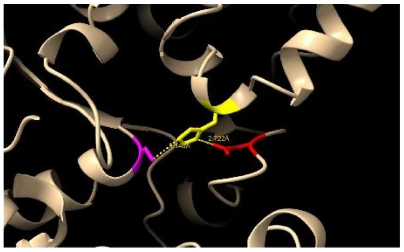

S98 H304 D199

Interatomic distances 3.140 and 2.722 Å. The pLDDT are 97.25, 92.62, and 90.75, respectively.
Model: AF-A0A2V2VWL8-F1-v6

**Unclassified Esterase (C4B63_35g372)**

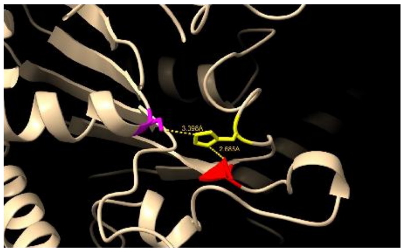

S211 H370 E336

Interatomic distances 3.398 and 2.685 Å (partially conserved). The pLDDT are 87.69, 83.50, and 84.81, respectively.
Model: AF-A0A2V2VED5-F1-v6

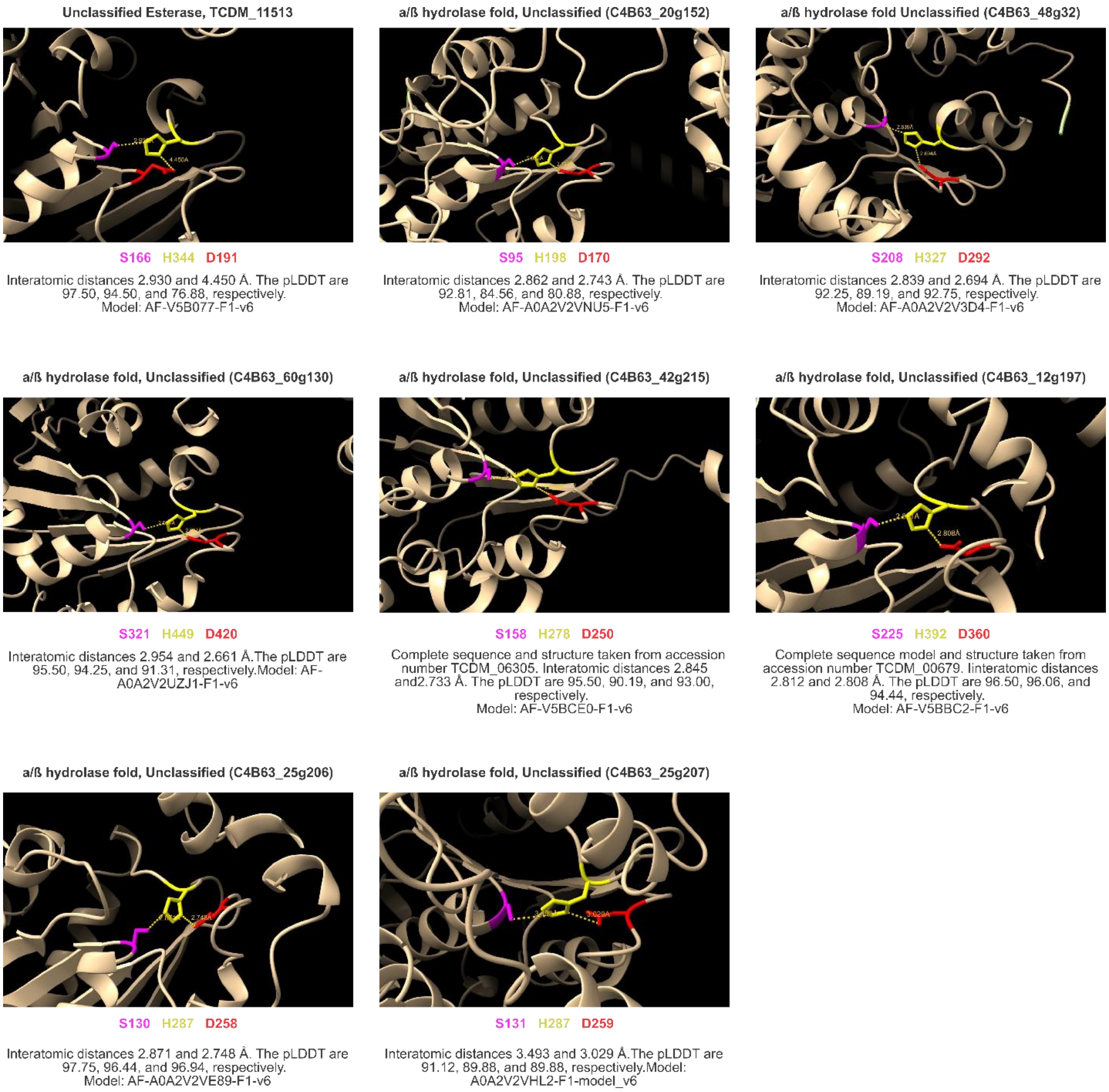

Structural analysis of the spatial arrangement of the catalytic Ser–His–Asp/Glu (S–H–D/E) triad for each protein**.** Residue numbers are indicated in the color-coded legend. Dashed lines denote interatomic distances measured in ångströms (Å). AlphaFold-predicted structures were obtained from the AlphaFold Protein Structure Database.

**Supplementary Figure S2**

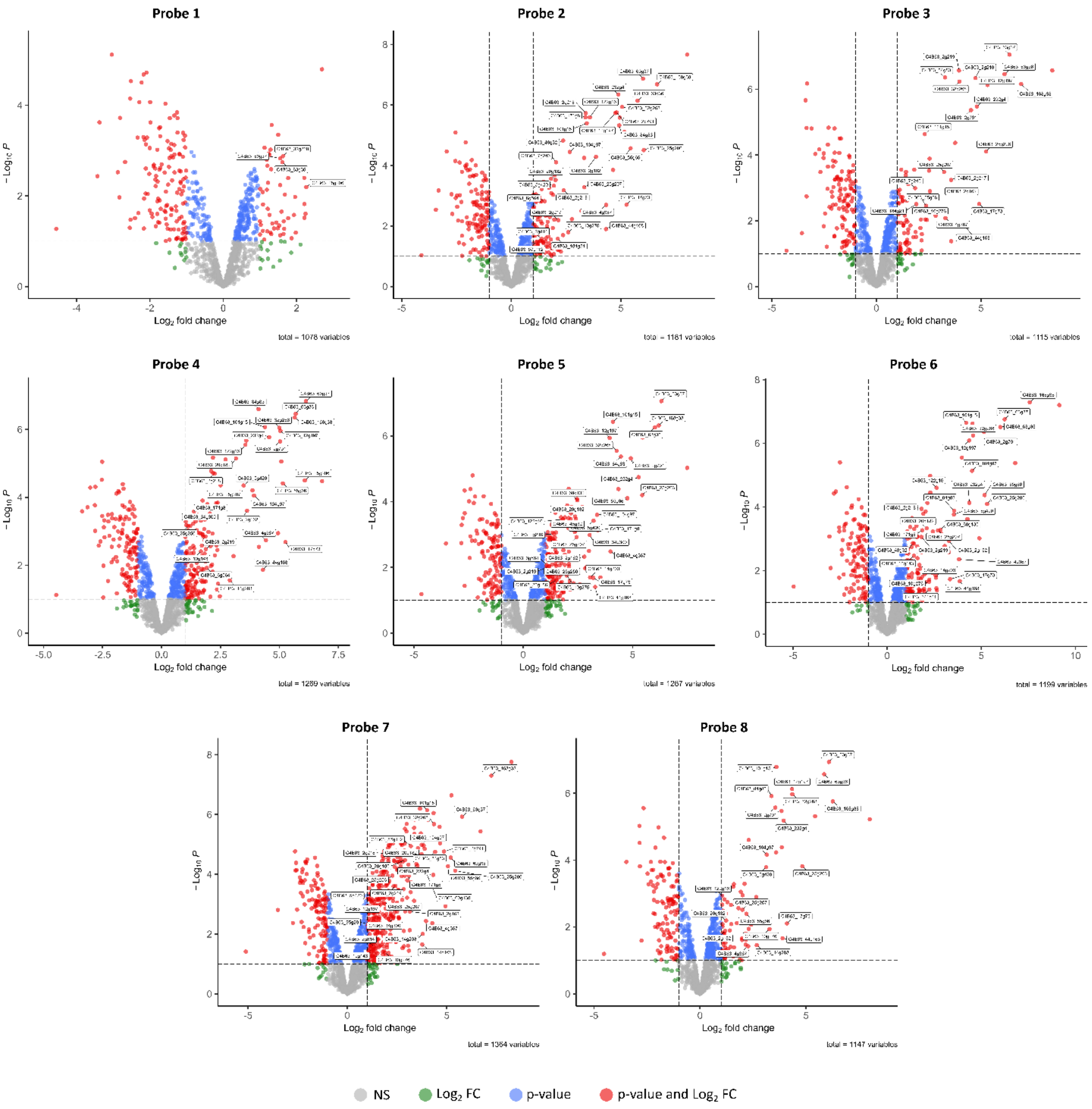

**Volcano plots of probe-enriched SHs in *T. cruzi***. Activity-based protein profiling (ABPP) coupled to label-free quantification mass spectrometry (LFQ-MS) was used to identify Serine Hydrolases (SHs) selectively enriched by nine activity-based probes (ABPs) in *T. cruzi* epimastigotes. The volcano plots shown are from the 2018 Dm28c assembly. Differential protein abundance between ABP-treated and DMSO control samples was assessed in biological replicates. In the volcano plots, the x-axis represents the $\log_2$ fold-change in enrichment, and the y-axis the $-\log_{10}$ p-value. Thresholds were set at $\log_2$ FC > 1 and P < 0.05. Proteins meeting both criteria are displayed on the right-hand side as red circles and correspond to *T. cruzi* SHs annotated by gene name. Green points indicate proteins with high fold-change but below-threshold significance, while blue points represent those with significant P values but lower fold-change. The complete list of curated SHs identified is provided in **Table 1**, while the complete list of proteins identified across all datasets is available in **Supplementary Data 3**.

#### Supplementary Figure S3

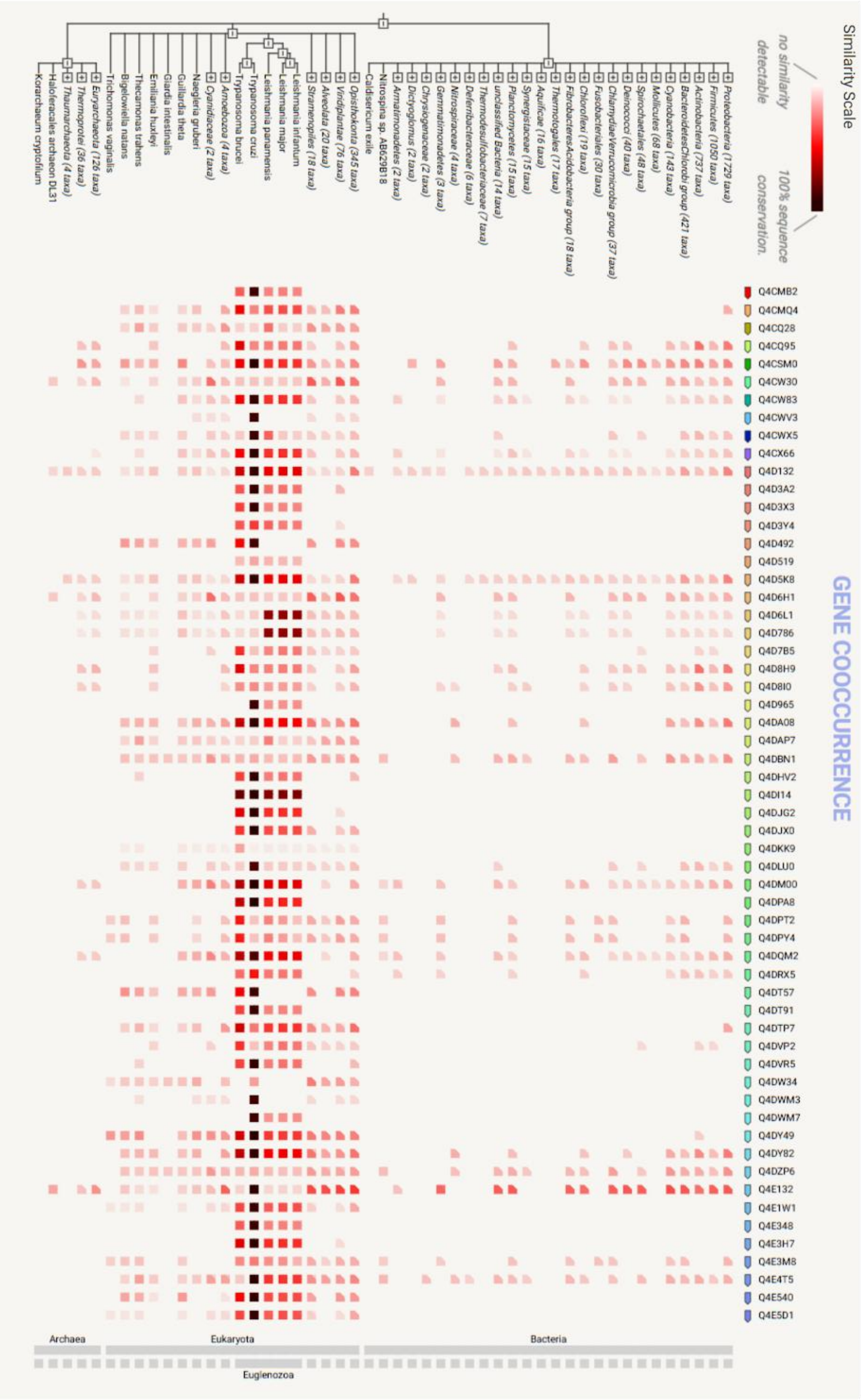

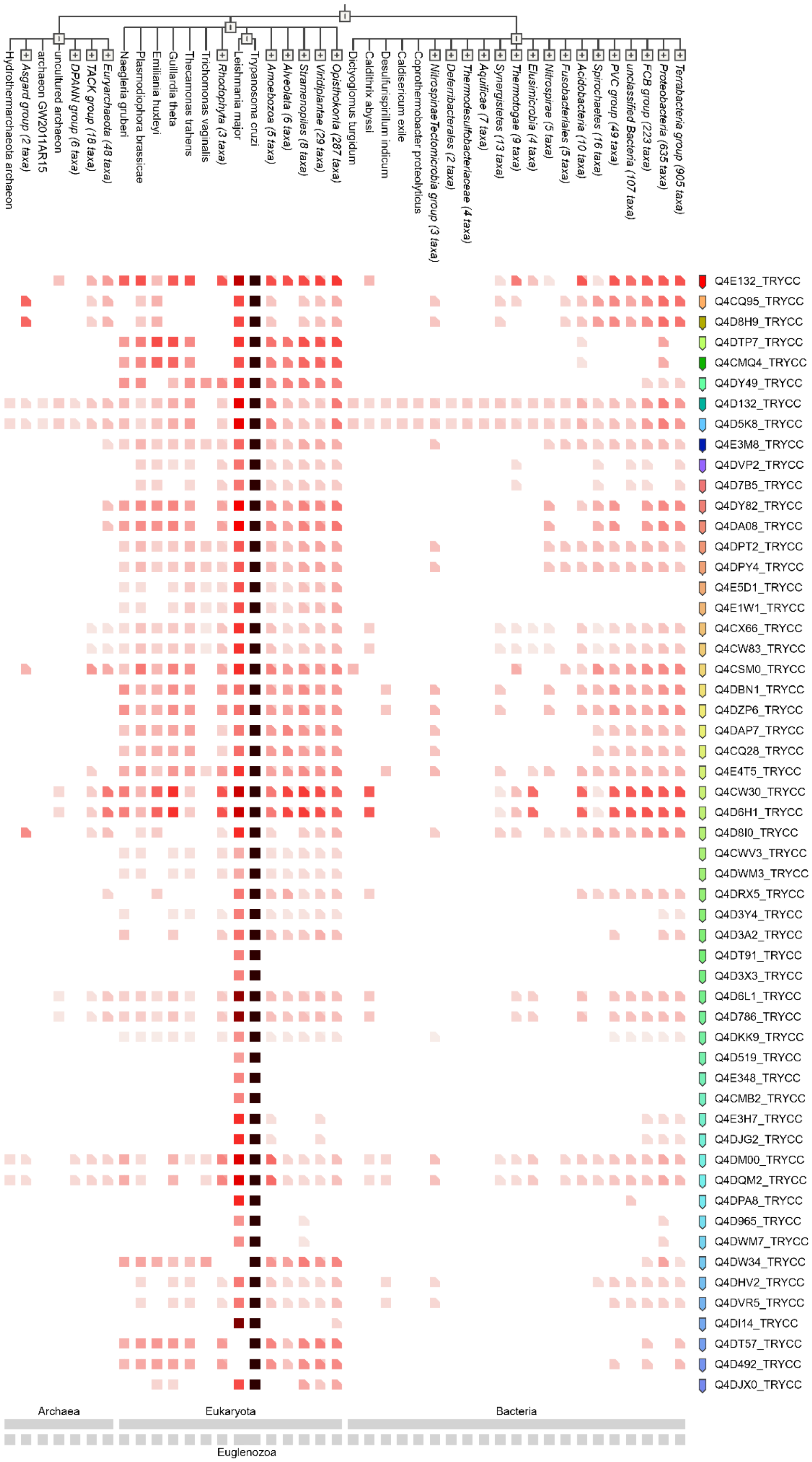
Hydrothermarchaeota archaeon
Asgard group (2 taxa)
archaeon GW2011AR15
uncultured archaeon
DPANN group (6 taxa)
TACK group (18 taxa)
Euryarchaeota (48 taxa)
Naegleria gruberi
Plasmodiophora brassicae
Emiliania huxleyi
Guillardia theta
Thecamonas trahens
Trichomonas vaginalis
Rhodophyta (3 taxa)
Leishmania major
Trypanosoma cruzi
Amoebozoa (5 taxa)
Alveolata (6 taxa)
Stramenopiles (8 taxa)
Viridiplantae (29 taxa)
Opisthokonta (287 taxa)
Dictyoglomus turgidum
Caldithrix abyssi
Desulfurispirillum indicum
Caldisericum exile
Coprothermobacter proteolyticus
Nitrospinae/Tectomicrobia group (3 taxa)
Deferribacterales (2 taxa)
Thermodesulfobacteriaceae (4 taxa)
Aquificae (7 taxa)
Synergistetes (13 taxa)
Thermotogae (9 taxa)
Elusimicrobia (4 taxa)
Nitrospirae (5 taxa)
Fusobacteriales (5 taxa)
Acidobacteria (10 taxa)
Spirochaetes (16 taxa)
PVC group (49 taxa)
unclassified Bacteria (107 taxa)
FCB group (223 taxa)
Proteobacteria (635 taxa)
Terrabacteria group (905 taxa)
Q4E132_TRYCC
Q4CQ95_TRYCC
Q4D8H9_TRYCC
Q4DTP7_TRYCC
Q4CMQ4_TRYCC
Q4DY49_TRYCC
Q4D132_TRYCC
Q4D5K8_TRYCC
Q4E3M8_TRYCC
Q4DVP2_TRYCC
Q4D7B5_TRYCC
Q4DY82_TRYCC
Q4DA08_TRYCC
Q4DPT2_TRYCC
Q4DPY4_TRYCC
Q4E5D1_TRYCC
Q4E1W1_TRYCC
Q4CX66_TRYCC
Q4CW83_TRYCC
Q4CSM0_TRYCC
Q4DBN1_TRYCC
Q4DZP6_TRYCC
Q4DAP7_TRYCC
Q4CQ28_TRYCC
Q4E4T5_TRYCC
Q4CW30_TRYCC
Q4D6H1_TRYCC
Q4D8I0_TRYCC
Q4CWV3_TRYCC
Q4DWM3_TRYCC
Q4DRX5_TRYCC
Q4D3Y4_TRYCC
Q4D3A2_TRYCC
Q4DT91_TRYCC
Q4D3X3_TRYCC
Q4D6L1_TRYCC
Q4D786_TRYCC
Q4DKK9_TRYCC
Q4D519_TRYCC
Q4E348_TRYCC
Q4CMB2_TRYCC
Q4E3H7_TRYCC
Q4DJG2_TRYCC
Q4DM00_TRYCC
Q4DQM2_TRYCC
Q4DPA8_TRYCC
Q4D965_TRYCC
Q4DWM7_TRYCC
Q4DW34_TRYCC
Q4DHV2_TRYCC
Q4DVR5_TRYCC
Q4DI14_TRYCC
Q4DT57_TRYCC
Q4D492_TRYCC
Q4DJX0_TRYCC
Archaea
Eukaryota
Bacteria
Euglenozoa

Co-occurrence of identified SHs across representative genomes from the three kingdoms of life. The colour denotes the similarity of its best hit in a given genome for each gene. All analyses were done using String v11.5 and v12 respectively.

#### METHODS

##### *Trypanosoma cruzi* cell culture

Dm28c epimastigotes were routinely maintained at mid-log growth phase by frequent dilution in LIT (5 g/L liver infusion broth, 5 g/L bacto-tryptose, 68 mM NaCl, 5.3 mM KCl, 22 mM $Na_2HPO_4$ and 0.4 % (w/v) glucose, pH 7.4) medium supplemented with 5 µM hemin and 10 % Foetal Bovine Serum (FBS), in static culture at 28 °C.

##### Protein extraction from *T. cruzi* parasites

Cells were collected by centrifugation at 3,500 × *g* for 5 minutes at 4 °C, washed three times with ice-cold Dulbecco's phosphate-buffered saline (PBS, pH 7.4), and lysed in buffer containing 25 mM Tris–HCl (pH 7.4), 150 mM NaCl, 1% Triton X-100, and 5% glycerol, supplemented with cOmplete™ Mini EDTA-free protease inhibitor cocktail (Roche, Cat. No. 1183170001). The lysates were then cleared by centrifugation at 13,000 × *g* for 10 minutes at 4 °C to remove insoluble debris. Protein concentrations were determined using the Pierce™ Rapid Gold BCA Protein Assay Kit (Thermo Fisher Scientific), following the manufacturer's instructions.

##### Live parasites ABP labelling

One hundred and fifty million *T. cruzi* epimastigotes (50 x $10^6$ epimastigote/mL) were incubated with the stated ABPs (10 µM) for 30 minutes at 28 °C, followed by lysis using the previously described procedure.

##### Cu-Catalysed azide-alkyne cycloaddition (CuAAC)

Biotin-$N_3$ (50 µM), $CuSO_4$ (1 mM), TBTA (0.1 mM), and sodium ascorbate (1 mM) were added to the ABP-labelled cell lysate (2 mg/mL). The click reaction mixture was incubated at room temperature for 1 hour with occasional mixing.

**Protein precipitation**

Following biotinylation via CuAAC, proteins were precipitated by adding nine volumes of ice-cold methanol and incubated overnight at –80 °C. The next day, samples were centrifuged at 10,000 × *g* for 10 minutes at 4 °C, and the resulting pellets were washed twice with ice-cold methanol to remove excess activity-based probes (ABPs), biotin-$N_3$, and residual click reagents. Pellets were then air-dried for 30 minutes before further processing.

**Affinity enrichment**

Precipitated proteins were resuspended in a minimal volume of 2% SDS in PBS and subsequently diluted to a final SDS concentration of 0.1%. NeutrAvidin–Agarose beads (50 µL per sample), pre-washed three times with four volumes of 0.1% SDS in PBS, were added to each sample. The mixtures were incubated for 2 hours at room temperature on an end-over-end rotating shaker to allow binding. Beads were then sequentially washed three times with 0.5% SDS in PBS, three times with 6 M urea in PBS, three times with PBS, and once with 50 mM triethylammonium bicarbonate (TEAB) buffer. Each wash was performed using 500 µL of the corresponding buffer, followed by centrifugation at 1,500 × *g* for 2 minutes at room temperature.

**On-bead reduction, alkylation, and tryptic digestion**

Beads obtained from the previous affinity enrichment step were incubated with 200 µL of 10 mM TCEP in 50 mM triethylammonium bicarbonate (TEAB) buffer for 45 minutes at 30 °C to reduce disulfide bonds. Following reduction, samples were washed with 400 µL of 50 mM TEAB, centrifuged at 1,500 × *g* for 2 minutes, and the supernatant discarded. The beads were then resuspended in 200 µL of 15 mM α-iodoacetamide in 50 mM TEAB and incubated in the dark for 45 minutes to alkylate free thiols. After alkylation, the beads were washed again with TEAB, resuspended in 200 µL of fresh 100 mM TEAB, and digested overnight with 4 µg of sequencing-grade modified trypsin at 37 °C for 16 hours. Following digestion, samples were centrifuged at 5,000 × *g* for 5 minutes, and the supernatants were collected. The remaining beads were washed twice with 50 µL of 50% acetonitrile (ACN) containing 0.1% formic acid (FA), centrifuged at 1,500 × *g* for 2 minutes, and the wash supernatants were pooled with the initial peptide eluate. The combined tryptic peptides were acidified to pH 3 using formic acid and dried by vacuum centrifugation. Peptides were then reconstituted in 0.1% (v/v) formic acid in water and desalted using Pierce™ Peptide Desalting Spin Columns (Thermo Scientific, Cat. No. 89851) according to the manufacturer's instructions. Finally, the eluates were dried completely under vacuum for downstream analysis.

**LC-MS analysis**

Peptide separations were carried out on a nanoHPLC Ultimate3000 (Thermo Scientific) using a nano column EASY-Spray ES903 (50 cm × 50 µm ID, PepMap RSLC C18). The mobile phase flow rate was 300 nL/min using 0.1 % formic acid in water (solvent A) and 0.1 % formic acid in acetonitrile (solvent B). The gradient profile was set as follows: 4-30 % solvent B for 114 min, 30-80 % solvent B for 14 min and 80 % solvent B for 2 min. MS analysis was performed using a Q-Exactive HF mass spectrometer (Thermo Scientific). For ionization, 1.9 kV of liquid junction voltage and 300 °C of capillary temperature were used. The full scan method employed a m/z 375–2,000 mass selection, an Orbitrap resolution of 120,000 (at m/z 200), a target automatic gain control (AGC) value of 1e6 and a maximum injection time of 100 ms. After the survey scan, the 8 most intense precursor ions were selected for MS/MS fragmentation. Fragmentation was performed with a normalized collision energy of 28 eV and MS/MS scans were acquired with a dynamic first mass. The AGC target was 5e5, resolution of 30,000 (at m/z 200), intensity threshold of 1.5e5, isolation window of 1.4 m/z units and maximum injection time of 55 ms. Charge state screening was enabled to reject unassigned, singly charged, and equal or more than seven protonated ions. A dynamic exclusion time of 30 s was used to discriminate against previously selected ions.

**Proteomics MS data processing**

All raw LC-MS/MS data were processed using FragPipe (v22.0).[1] Protein identification was carried out using MSFragger (v4.1),[2] and label-free quantification was performed using IonQuant (v1.10.27).[3] Searches were performed as two independent FragPipe runs, each against a separate *T. cruzi* Dm28c genome assembly (2018 and 2014). The following search parameters were used: trypsin digestion with maximum 2 missed cleavages, carbamidomethylation of cysteine as a fixed modification, oxidation of methionine, acetylation of protein N-termini as variable modifications, minimum peptide length of 7, a maximum number of modifications per peptide set at 3, and protein false discovery rate (FDR) 0.01. All subsequent data processing and statistical analysis were performed in R (v4.5.0), using scripts adapted by the authors from open-source scripts by Richter and Isern et al..[4] LFQ intensity values were $\log_2$-transformed to normalize distribution. Data distribution and variability were assessed using density plots (*ggplot2*),[5] boxplots, and mean-standard deviation plots (*IceR*). Missing data were imputed using a left-censored normal distribution approach, akin to the Perseus strategy.[6] Specifically, missing values were replaced with random values drawn from a normal distribution shifted 1.8 standard deviations below the mean and scaled by a factor of 0.3. The quality of the imputation was assessed via replicate correlation plots and distributional comparisons before and after imputation. Differential abundance analysis was conducted using the *limma* package.[7] A linear model was fitted for each protein across the two conditions (e.g., “Probe” vs. “Ctrl”), empirical Bayes moderation was applied to the variance estimates. Proteins were considered differentially abundant based on a dual threshold of $|\log_2(\text{fold change})| > 1$ and $p\text{-value} < 0.05$. Although

Benjamini–Hochberg false discovery rate (FDR)-adjusted *p*-values were computed, nominal p-values were used for thresholding given the target-discovery nature of this workflow, so as not to discard genuine low-abundance targets; candidate hits were subsequently prioritized by manual curation. Protein identifiers were cleaned and standardized to ensure uniqueness prior to analysis. Each run was analysed separately, and the serine hydrolases identified as enriched across the two assembly searches were combined into a single set of candidate targets. Significant protein abundance changes were visualized using *EnhancedVolcano* (version 1.14.0).

***In silico* Identification of Serine hydrolases in *T. cruzi* genome**

Predicted protein sequences from 2018 genome strain Dm28c 2018 of *T. cruzi* were downloaded from TriTrypDB (Assembly GCA_003177105.1, May 30, 2018).[8] Predicted serine peptidases (44 accession numbers) were identified via a BLASTP search (BLAST+ 2.12.0) against the MEROPS Scan Sequences database (file "merops_scan.lib", MEROPS release 12.5, 8 September 2023; downloaded 2026-02),[9] using default parameters and a threshold E-value of 1 x $10^{-5}$. Because the conserved peptidase domain, and not the full-length protein, is used as the query seed, query-coverage thresholds were not applicable; selection was based on the E-value rather than percent identity, which does not carry comparable statistical meaning. Predicted serine hydrolases (105 accession numbers) were identified using hmmscan (HMMER 3.4) with Pfam-A HMM models (downloaded from https://ftp.ebi.ac.uk/pub/databases/Pfam/current_release/; released 2026-01-13 13:49), selecting the Pfam families assigned to serine hydrolases with a threshold on the independent-domain E-value (i-E-value) of 1 x $10^{-5}$. If several Pfam domains are detected, they appear sorted by increasing E-value. The serine-hydrolase Pfam families used were PF00326, PF00450, PF00561, PF00930, PF01425, PF01694, PF01734, PF01764, PF02230, PF02897, PF03403, PF05577, PF07819, PF12146, PF12697, PF00082, PF00089, PF00717, PF01343, PF02129, PF03572, PF03575, PF04096, PF10502, PF13365, PF07859, PF00657, PF00756,

PF02450, PF03959, PF05057, PF08538, PF10230, PF10503, PF13472, PF00975, PF01674, PF02253, PF00151, PF20434, PF08386, PF03096, PF11339 and PF12695 (**Supplementary Data 1**). In addition, we also selected 31 accession numbers with InterPro, CDD or Panther domains as serine hydrolases in TriTrypDB,[10] but not selected previously by Pfam domains. A total of 135 unique proteins were identified and they are shown in Table ; the identification route(s) (MEROPS, Pfam and/or InterPro/CDD/Panther) by which each gene was detected are indicated in Supplementary Data 1 Identical proteins (≥ 95% identity) were grouped as "Other copies" using CD-HIT 4.8.1 (-c 0.95, -n 5) (42 proteins).[11] Proteins with 90-99% identity but with smaller length, were grouped as "Fragmented copies" (13 proteins). In the case of C4B63_104g97, 8 accession numbers with 90-95% identity and the same sequence length were grouped in the same cluster. The accession numbers C4B63_4g438, C4B63_38g92, C4B63_42g174, and C4B63_113g62 with PDZ domains were discarded because their sequences do not have a serine hydrolase domain. The accession numbers C4B63_128g64, C4B63_128g65, and C4B63_128g66 were also discarded since they are fragments of the same gene, but the complete sequence was not found in any of the *T. cruzi* genomes. The sequence of the accession number C4B63_63g36 is a fragment of the same gene annotated in C4B63_63g37. The list was reduced to 73 unique proteins and the product description was taken from TriTrypDB. Protein length was calculated from the primary sequence and validated by comparison with other *T. cruzi* strains. Any corrected lengths are provided in parentheses in Table S1. We used the corrected sequence in the further analysis (CURATION AND CLASSIFICATION). To verify that each protein is a Serine Hydrolase, the nucleophilic serine in the motif GxSxG and the catalytic triad or dyad were searched in the protein sequence. We identified the catalytic triad or dyad by BLASTP with proteins of MEROPS Scan Sequences database and/or in the alignments done by CDD-search[12] or Hmmer-scan.[13] We then assessed whether the catalytic residues were geometrically close in the AlphaFold predicted structure, retrieved from the AlphaFold Protein Structure Database (https://alphafold.ebi.ac.uk) linked to TriTrypDB; where the Dm28c 2018 sequence was truncated, the corrected sequence was used.[14,15] Catalytic geometry was scored as conserved when the Ser Oγ–His Nε2 and His–Asp/Glu interatomic distances were within 5 Å and the participating residues had an average pLDDT ≥ 80 (a few cases with pLDDT between 60 and 79 were also accepted). The type of triad or dyad were searched according to the classification of the enzyme. The proteins were classified as esterase, serine peptidase, or amidase according to Pfam, product description, MEROPS classification, other identified domains, or catalytic triad. BLASTP was performed against protein databases of other organisms to verify if these proteins belong to the peptidase, lipase, or esterase families in cases of conflicting classifications. Also, the esterases and peptidases were classified into MEROPS families or subclassification when possible. In families S33 and S9, there are a large number of non-peptidase homologues. The family S9 and S33 have similar organization of the catalytic triad (S-D-H), but peptidases S9 have an additional domain - a beta-propeller domain. Also, S15 family has an extra domain: a C-terminal β-sandwich domain.[16] "Additional information" column include Pfam with E-value higher than $1 \times 10^{-5}$, BLASTP additional searches, the compared catalytic triad, and/or information about protein fragments. From the 73 unique proteins, 56 were selected as having a conserved or partially conserved catalytic triad (**Table 1** and **Supplementary Fig. S1**); the criteria used to define a conserved or partially conserved triad are detailed in **Supplementary Data 1**.

##### ***In silico*** **analysis of domains in enriched proteins**

Pfam domains and transmembrane regions were obtained from https://www.ebi.ac.uk/interpro/[17] and Pfam protein families database.[18] Gene accession numbers belonging to *Leishmania mexicana* strain MHOM/GT/2001/U1103 genome (assembly GCA_000234665.4) and gene accession numbers belonging to *Trypanosoma brucei brucei* TREU927 (assembly GCA_000002445.1) were obtained from TriTrypDB. Localization of *T. brucei* genes was obtained from experimental data in TrypTagDB.[19] For Proteins localized to organelles, only C-terminal tagging is considered, except for C4B63_32g268 that was detected in cytosol and glycosomes in *T. brucei*.[20]

#### iBAQ Analysis and Protein Ranking

iBAQ (intensity-based absolute quantification) analysis was performed to estimate relative protein abundances across all experimental conditions. ProteinGroups.tsv output from FragPipe (v 22.0)[1] was processed in R (v4.5.0), using scripts adapted by the authors from open-source scripts by Richter and Isern et al..[4] Sample-wise intensity values were extracted based on a curated annotation file and log-transformed. Proteins annotated as contaminants were excluded. To calculate iBAQ values, in silico tryptic digestion of the 2018 reference proteome of strain Dm28c of *T. cruzi* was performed using the Biostrings R package,[21] applying canonical cleavage after K/R not followed by P, and filtering peptides to a length range of 6–30 amino acids. The number of theoretically observable tryptic peptides per protein was computed and used to normalize the summed MS1 intensities across all replicates, yielding iBAQ values. Proteins with zero or missing intensities or lacking theoretical peptides were excluded. Remaining proteins were ranked by decreasing iBAQ, and their $\log_{10}$-transformed values plotted against rank. A panel of target genes of interest was highlighted within the distribution using the ggplot2 and ggrepel packages.
